# Rapid characterization of spike variants via mammalian cell surface display

**DOI:** 10.1101/2021.03.30.437622

**Authors:** Kamyab Javanmardi, Chia-Wei Chou, Cynthia I. Terrace, Ankur Annapareddy, Tamer S. Kaoud, Qingqing Guo, Josh Lutgens, Hayley Zorkic, Andrew P. Horton, Elizabeth C. Gardner, Giaochau Nguyen, Daniel R. Boutz, Jule Goike, William N. Voss, Hung-Che Kuo, Kevin N Dalby, Jimmy D. Gollihar, Ilya J. Finkelstein

## Abstract

The SARS-CoV-2 spike (S) protein is a critical component of subunit vaccines and a target for neutralizing antibodies. Spike is also undergoing immunogenic selection with clinical variants that increase infectivity and partially escape convalescent plasma. Here, we describe spike display, a high-throughput platform to rapidly characterize glycosylated spike ectodomains across multiple coronavirus-family proteins. We assayed ∼200 variant SARS-CoV-2 spikes for their expression, ACE2 binding, and recognition by thirteen neutralizing antibodies (nAbs). An alanine scan of all five N-terminal domain (NTD) loops highlights a public class of epitopes in the N1, N3, and N5 loops that are recognized by most of the NTD-binding nAbs. Some clinical NTD substitutions abrogate binding to these epitopes but are circulating at low frequencies around the globe. NTD mutations in variants of concern B.1.1.7 (United Kingdom), B.1.351 (South Africa), B.1.1.248 (Brazil), and B.1.427/B.1.429 (California) impact spike expression and escape most NTD-targeting nAbs. However, two classes of NTD nAbs still bind B.1.1.7 spikes and neutralize in pseudoviral assays. B.1.351 and B.1.1.248 include compensatory mutations that either increase spike expression or increase ACE2 binding affinity. Finally, B.1.351 and B.1.1.248 completely escape a potent ACE2 peptide mimic. We anticipate that spike display will accelerate antigen design, deep scanning mutagenesis, and antibody epitope mapping for SARS-CoV-2 and other emerging viral threats.

## Introduction

Severe acute respiratory syndrome coronavirus 2 (SARS-CoV-2) is the causative agent of the COVID-19 pandemic, causing >127 million infections and >2.7 million deaths worldwide (as of 28/Mar/2021). Related betacoronaviruses SARS-CoV-1 and Middle Eastern Respiratory Syndrome (MERS) have also caused epidemics in 2002 and 2012, respectively^1–3^. Human coronavirus HKU1, first discovered in 2004, often manifests as a mild upper respiratory disease^4^. The large reservoir of diverse and endemic coronaviruses in animals, and their frequent zoonotic transmission, suggests that future human outbreaks are inevitable^5–7^.

Coronaviruses infect cells via attachment of viral transmembrane spike (S) glycoproteins^8^. SARS-CoV-2 spike interacts with angiotensin-converting enzyme 2 (ACE2) and other cell surface receptors to mediate fusion between the virus envelope and cell membrane^9–11^. Spike homotrimers consist of the S1 and S2 functional subdomains^12^. After spike binds ACE2, structural rearrangements in the spike and cleavage by host proteases separate the S1 subunit from the S2 stalk^8^. The S2 stalk then undergoes further conformational changes that lead to membrane fusion and cell entry. The S1 subunit, which is composed of the N-terminal domain (NTD) and receptor-binding domain (RBD)^12^, is the key determinant of tissue and host tropism^8^.

Humoral immunity to the spike glycoprotein is the most potent means of protection from SARS-CoV-2^13^. SARS-CoV-2 vaccines generate a strong polyclonal antibody response by delivering spikes via immunization^14–16^. Spike is also the primary target for prophylactic and therapeutic neutralizing monoclonal antibodies (nAbs) and ACE2 binding inhibitors^17–21^. However, spike continues to mutate and recombine, establishing new variants for immunogenic selection^22^. Multiple variants of concern (VOCs) have increased viral transmissibility and antibody escape^6,22^. Since the emergence of a globally dominant D614G mutation^23–25^, newer VOCs with compound spike mutations have taken hold. The B.1.1.7 (United Kingdom)^26,27^, B.1.351 (South Africa)^28^, B.1.1.248 (Brazil)^29^, and B.1.427/B.1.429 (California)^30^ lineages are of particular concern because they partially evade monoclonal antibodies, convalescent sera, and vaccine-induced humoral immunity^31^. The antigenicity and infectivity of new virus variants are often assayed via live virus, pseudotyped virus, and animal protection experiments. These assays are low throughput and require lengthy viral preparations^28^.

Cell surface display of the spike protein or its sub-domains is a high-throughput platform approach to functionally characterize key aspects of SARS-CoV-2 variants^32,33^. For example, Starr et. al. and others have expressed the RBD on the surface of yeast cells to measure the sub-domain expression, ACE2 binding affinity, and RBD-targeting nAb escape^20,34–37^. However, all spike VOCs include critical mutations that are outside the RBD. Moreover, the humoral immune response produces potent neutralizing antibodies that target the NTD as well as the RBD. Here, we describe a new experimental platform that measures spike expression, receptor binding, and antibody escape across variant spike homotrimers on the surface of mammalian cells.

Spike display is a high-throughput platform to characterize spike glycoproteins from diverse coronavirus families. Complex spike variants are displayed on the surface of mammalian cells and assayed via flow cytometry. Spikes can be cleaved from cell surfaces to further accelerate structural and biophysical characterization. Using this platform, we mapped an NTD supersite that is recognized by the majority of NTD-directed nAbs. We also characterized the individual mutations composing the B.1.1.7, B.1.351, B.1.1.248, and B.1.427/B.1.429 variants using flow cytometry and biolayer interferometry (BLI). All four complex variants show escape from NTD-targeting nAbs while B.1.351, B.1.1248, and to a lesser degree B.1.427/B.1.429, escape some RBD-targeting nAbs. Destabilizing mutations such as Δ242-244 and R246I in B.1.351, which enable nAb escape, are compensated for by stabilizing mutations D215G and K417N. The conserved N501Y mutation, found in B.1.17, B.1.351, and B.1.1.248 also increases ACE2 binding affinity. Most VOCs also escape the ACE2-mimetic LCB1 peptide, suggesting that micropeptide inhibitors must be updated or used as part of a multi-component inhibitor cocktail^17^. We anticipate that spike display will accelerate antigen design, spike characterization, and epitope mapping to aid ongoing and future pandemic countermeasures.

## Results

### Assessing spike variants on mammalian cell surfaces

We express the SARS-CoV-2 spike ectodomain on the surface of human embryonic kidney (HEK293T) cells. Six proline substitutions stabilize the homotrimeric complex in the pre-fusion state, along with the globally dominant D614G mutation (termed 6P-D614G)^23,25,38^. Spike is directed to cell membranes via an N-terminal Ig Kappa secretion signal and a C-terminal PDGFR-β transmembrane domain^39^. The 58 amino acid (aa) flexible linker includes a triple FLAG (3xFLAG) epitope tag as a proxy for expression, a StrepII tag for purification, and a 3C protease cleavage site (Fig. 1a). Immunostained fixed cells show spike at the cell membrane (Fig. 1b and Supplementary Fig. 1). We confirmed the native homotrimeric assembly via negative-stain electron microscopy (nsEM) of spikes that were cleaved from the cell surface by 3C protease (Fig. 1c). Two-dimensional class averages indicate that >99% of the surface-displayed spikes are in the pre-fusion conformation, with the majority (71%) of the particles having one receptor-binding domain (RBDs) up (Supplementary Fig. 3). These results are consistent with previous assessments of HexaPro (6P) and D614G spike RBD up-down equilibrium^12,24,38^.

**Figure 1:**
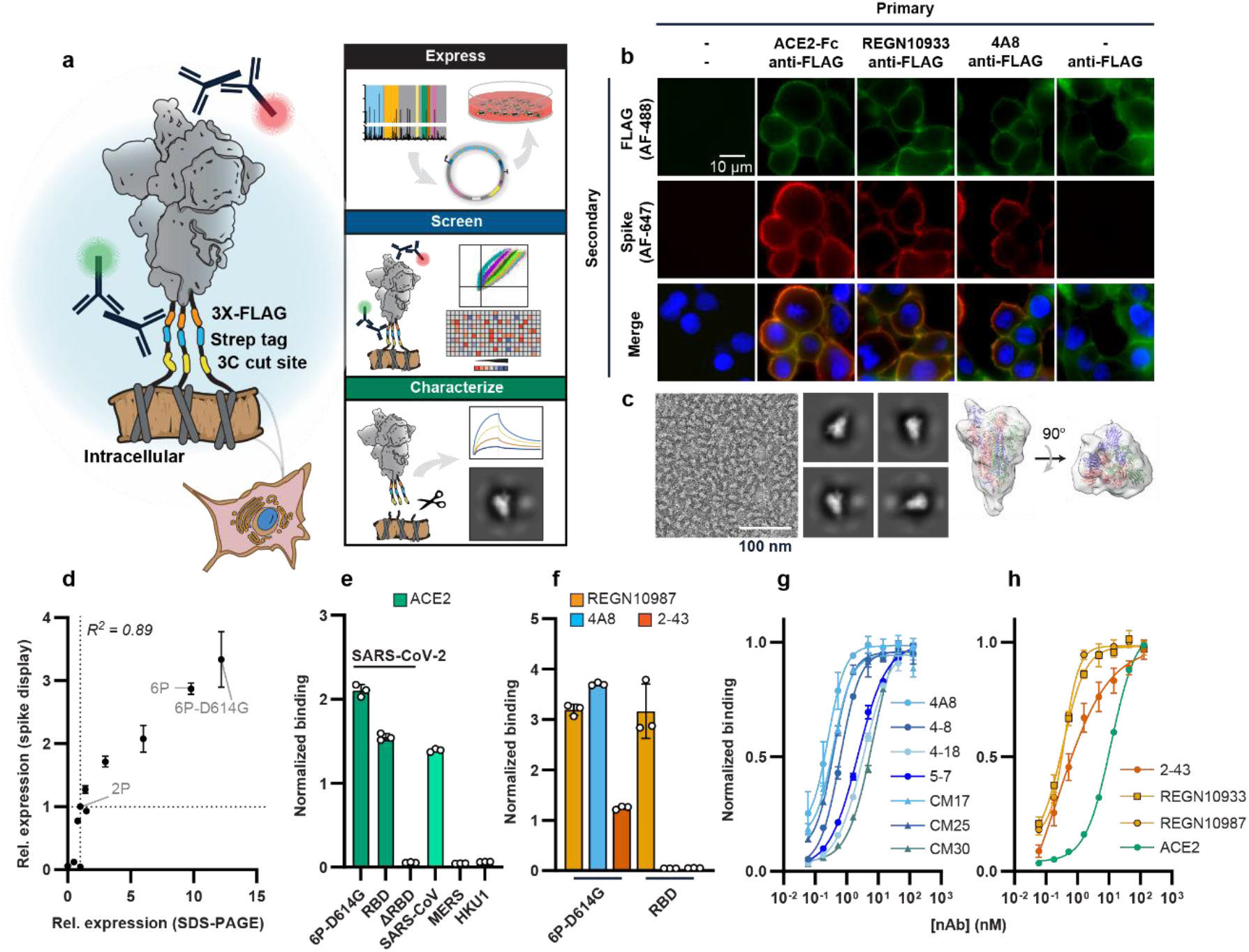
Biophysical characterization of spikes displayed on human cells. **(a)** Spike ectodomains are displayed on the surface of HEK293T cells. An automated cloning pipeline is coupled with flow cytometry to enable high throughput screening. Biophysical characterization is performed with spikes cleaved from cell surfaces. **(b)** Immunostaining confirms that SARS-CoV-2 (6P-D614G) spikes are localized to cell membranes and bind ACE2 (RBD-directed), REGN10933 (RBD-directed), or 4A8 (NTD-directed). Scale bar: 10 µm. **(c)** Negative-stain electron microscopy micrograph (left), 2D-class averages (middle), and a 3D model of surface-displayed spikes in the pre-fusion conformation. The majority of particles show 1-RBD up configuration. Scale bar: 100 nm **(d)** Relative spike display signal correlates with recombinant spike expression levels for engineered and clinical spike variants^24,37^. For both axes, the signal is normalized to spike-2P expression. Pearson correlation is used for statistical analysis. **(e)** ACE2 soluble domain binding by diverse coronavirus-family spikes displayed on mammalian cell surfaces. **(f)** Normalized binding for RBD- (REGN10987), NTD- (4A8), and S1-directed (2-43) neutralizing antibodies was measured using either the full SARS-CoV-2 spike or the isolated RBD. **(g)** Titration of NTD binding nAbs using spike display and flow cytometry. **(h)** Titration of ACE2, RBD-directed (REGN10933 and REGN10987), and S1-directed (2-43) nAbs using spike display and flow cytometry. All measurements in (E-H) are an average of three biological replicates. Error bars: S.D.

We assessed spike expression and antigenicity via flow cytometry. SARS-CoV-2 spike variants and related spike homologs were rapidly constructed via Golden Gate assembly from standardized parts and acoustic liquid handling (Fig. 1a and Supplementary Fig. 2 and Methods). Each variant was transfected into HEK293T cells and stained with anti-FLAG and fluorescent secondary (anti-mouse) antibodies. Mutant SARS-CoV-2 spike expression on the surface of mammalian cells strongly correlated with the relative expression of recombinant spike mutants (Fig. 1d)^25,38^. The relative expression of spikes from related betacoronaviruses SARS-CoV-1, MERS, and HKU1 also followed the relative stabilities of their recombinant proteins (Supplementary Fig. 5a-b)^12,38,40,41^. We next tested whether spike display can be used for antigen design by testing pre-fusion stabilizing proline substitutions into SARS-CoV-1 spike (Supplementary Fig. 5d-e). Spike expression improved 5-fold, suggesting further stabilization over the parental SARS-CoV-1 construct. We conclude that spike display can be used for the rapid assessment of structural expression of coronavirus spike variants and for designing next-generation pre-fusion stabilized vaccine targets.

Next, we measured ACE2 binding affinity of surface-displayed spikes. Non-cross reactive fluorescent secondary antibodies were directed against the human (ACE2 binding) and mouse Fc (anti-FLAG; spike expression). This two-color assay enables spike expression-based signal normalization for antigen binding (Fig. 1e and Supplementary Fig. 4). As expected, SARS-CoV-1, SARS-CoV-2 and the isolated RBD avidly bound ACE2, whereas spike(ΔRBD) did not bind ACE2 above background. The higher affinity of SARS-CoV-2 relative to CoV-1 spike is consistent with prior *in vitro* measurements with purified proteins^12^. The MERS and HKU1 spikes recognize the human DPP4 receptor and 9-O-acetylated sialic acids respectively, and have no affinity for ACE2^42^.

Spike display enables rapid flow cytometry-based characterization of neutralizing antibodies (nAbs) ^32^. An estimated 10% of the nAbs in seroconverted COVID-19 patients bind outside of the RBD and many RBD-binders may bind across multiple subunits^18,32,43,44^. Therefore, we assayed nAb binding to different spike sub-domains. We first tested REGN10987, 4A8, and 2-43 which are RBD-, NTD-, and S1 (quaternary)-binding nAbs, respectively (Fig. 1f)^43–45^. The signal generated from human-Fc binding secondary antibodies was normalized to the spike expression signal, enabling accurate binding measurements (see Methods). We observed domain-specific nAb binding with full spikes, whereas the RBD alone only bound REGN10987. We next measured quantitative binding affinities of seven NTD-binding nAbs via surface display (Fig. 1g). We selected nAbs that bind with pM to ∼10 nM binding affinities. These peripheral blood mononuclear cell (PBMC) derived antibodies were identified through diverse discovery platforms, including Ig-seq^46^, and exhibited differing neutralization potencies and NTD binding epitopes. The binding affinities derived from fitting the sigmoidal titration curves corresponded closely to the affinities measured with recombinant proteins (Supplementary Fig. 6 and Table S1). Titration experiments with ACE2, the REGN-COV2 cocktail nAbs (REGN10933 and REGN10987, both binding the RBD), and 2-43 (S1 binder) confirmed that spike display is a quantitative platform for measuring antibody and antigen-binding affinities. (Fig. 1h). Assaying nAb binding on full-length spike ectodomains (S-ECD) will thus be a valuable tool for rapid antibody discovery and characterization.

### Immune escape from public epitopes in the spike N-terminal Domain

The SARS-CoV-2 spike NTD elicits high affinity neutralizing antibodies in seroconverted and vaccinated patients^43,44,47^. However, 46% of all spike protein mutations (399,299 out of 866,373 total; GISAID database accessed on 24/Feb/2021) are in the NTD—two-fold higher than a random distribution (expected to be 23%) (Fig. 2a, Supplementary Fig. 7, and Methods)^48^. These clinical NTD mutations raise the possibility of immune pressure and subsequent escape. To elucidate the mechanisms for binding and neutralization, we focused on ten neutralizing NTD-directed antibodies, six of which had their epitopes mapped via cryo-EM^43,44,47,49^. Notably, 4A8, CM25, and 4-8 bind loops N3 (residues 141 to 156) and N5 (residues 246-260), suggesting that these may be public epitopes^49,50^. We cloned 72 alanine spike mutants that encompass all five NTD loops and adjacent residues (Fig. 2b). Alanine NTD substitutions very mildly destabilized spike expression (Supplementary Fig. 8). We tested this alanine library against the ten NTD nAbs described above, along with the RBD-binding ACE2 and REGN10987 as negative controls (Fig. 2c).

**Figure 2:**
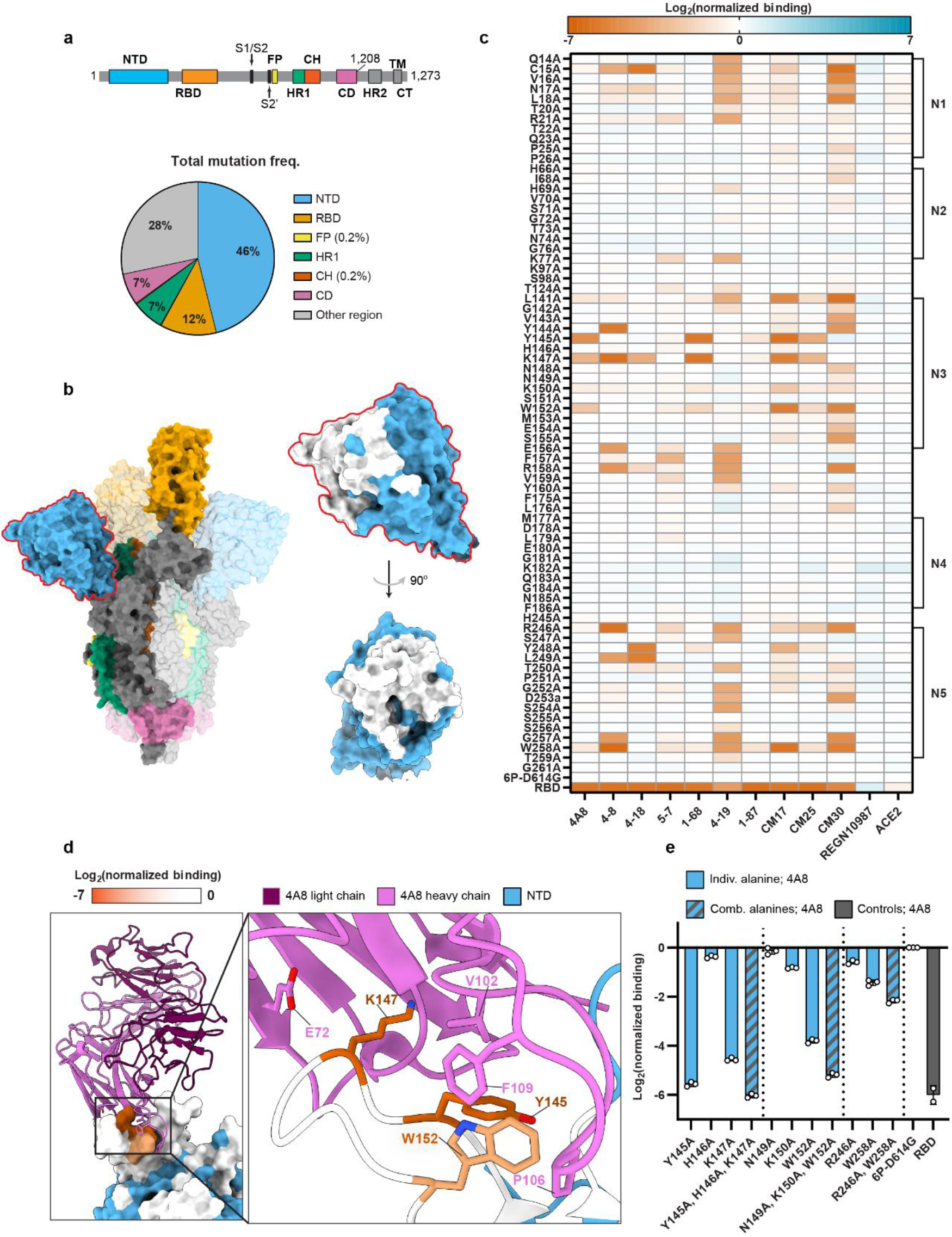
A high-resolution map of NTD-targeting nAb epitopes. **(a)** Spike domain map (top)^12^, and distribution of all non-synonymous mutations (total = 866,373) found in GISAID (accessed on 24/Feb/2021) (bottom) The NTD harbors 46% of all mutations while making up 23% of the protein. **(b)** Spike trimer structure (PDB: 7DDN^67^) with domains colored as in panel (a). An enlarged structure of the NTD (blue) with alanine scan positions (white) is shown on the right. **(c)** The effect of single alanine substitutions on antibody and ACE2 binding measured by flow cytometry (See Methods and File S1 for the mean and S.D.). Red = decreased binding, blue = increased binding relative to the reference spike (6P-D614G). The RBD is included as a negative control for all NTD-binding antibodies (last row). NTD loops 1-5 annotated on the right. **(d)** Co-structure (PDB: 7C2L^43^) of the 4A8 Fab (light chain = violet, heavy chain = pink) in complex with the NTD (blue). Alanine scan binding data for 4A8 is superimposed on the NTD and represented on a 0 to −7 binding scale. **(e)** Combining multiple alanine mutations abrogates 4A8 binding. RBD is included as a negative control (gray). Mean ± S.D. of at least two biological replicates.

We first focused on 4A8, which interacts with eight residues in the N3 and N5 loops of the NTD^43^. Unexpectedly, only three of these substitutions—Tyr145, Lys147, and Trp152—reduced antibody binding >5-fold (Fig. 2c and 2d). Combining any two substitutions resulted in an additional loss of binding, sometimes to below the detection limit (Fig. 2e). Surprisingly, two close contacts in the 4A8 Fab-spike ectodomain structure are dispensable for strong binding. Lys150, which forms a salt bridge with 4A8, only minimally reduced binding affinity. Arg246 on the N5 loop, which is buried in the 4A8-spike interface, reduced binding a modest 0.8-fold. Antibodies CM17, CM25, and 1-68 were also sensitive to alanine substitutions at Tyr145, Lys147, and Trp152, suggesting that they recognize the same NTD epitope as 4A8^47^. Despite competing with 1-68 in a spike binding assay, 1-87 showed little to no loss in binding from any single-alanine substitution in our library. We speculate that the CDR-1 and -3 of the 1-87 are minimally impacted by single alanine substitutions at the NTD surface. Antibodies 5-7 and 4-19 were more sensitive to substitutions in the N1 (residues 14-26) and N5 loops and a region between the N3 and N4 loops. Both of these nAbs have weaker neutralizing IC50’s (5-7 = 0.033 μg mL^−1^ and 4-19 = 0.109 μg mL^−1^) relative to nAbs which showed strong N3 loop binding (4-8 = 0.009 μg mL^−1^, 1-68 = 0.014 μg mL^−^1, CM25 = 0.012 μg mL^−1^ IC50s), suggesting that the N3 loop dominates NTD-mediated spike neutralization^44^. On the contrary, mutations in the N2 and N4 loops were inconsequential to antibody binding despite their proximity to the NTD supersite. Spike display complements structural studies and Fab competition assays by providing high-resolution epitope maps and by reporting affinity changes as a consequence of amino acid substitutions.

### Circulating spike mutants escape NTD- and RBD-directed antibodies

Guided by the alanine scan results, we profiled 46 non-synonymous NTD mutants for their impact on spike expression and antibody evasion. All NTD mutants retained wildtype levels of ACE2 binding and spike expression (Fig. 3a). Substitutions at Tyr145 strongly reduced 4A8 binding, as highlighted by the alanine scan. These polar and charged residues (Asn, Asp, and His) likely disrupt the hydrophobic and/or π/π interactions with 4A8’s CDR-3 region. Substitutions at Tyr145 also reduced CM17, CM25, and 1-68 binding. NTD variants that reduced nAb binding >16-fold appeared at very low frequencies (<0.001) in the GISAID database. Two noteworthy exceptions include M153T and S254F, both of which show strong escape from CM30 (Fig. 3b and Supplementary Fig. 9)^48^.

**Figure 3:**
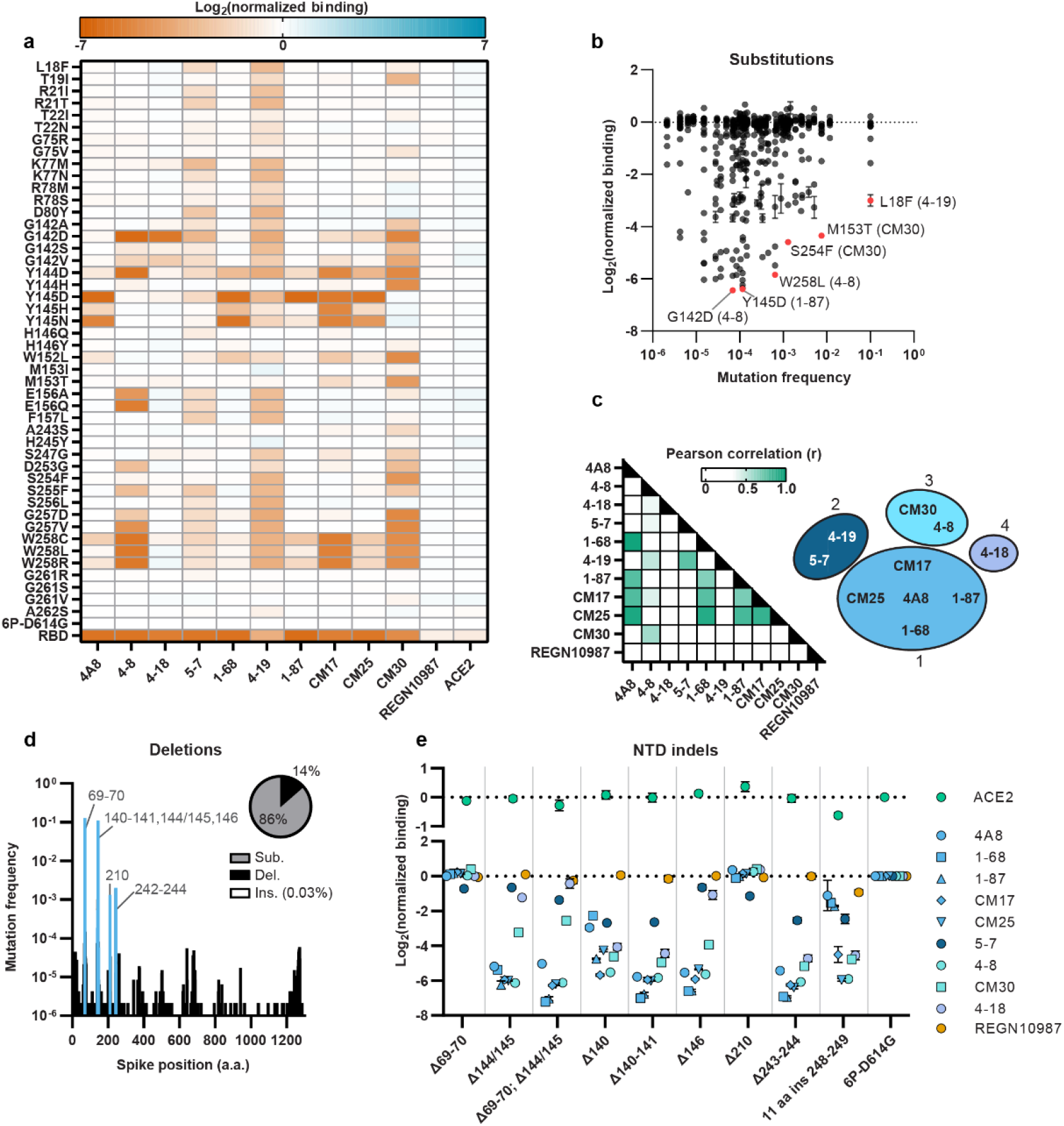
Clinical NTD mutants evade nAb binding. **(a)** The effect of single amino acid substitutions on antibody or ACE2 binding measured by flow cytometry. Red = decreased binding, blue = increased binding, relative to spike (6P-D614G). RBD is a negative control for all NTD binding antibodies (last row). **(b)** Nearly all nAb-evading NTD mutants occur at a low frequency in the GISAID. Notable mutations and the antibodies they evade are highlighted in red. **(c)** Pearson correlation matrix comparing the loss of binding across all antibodies and point mutants (left). The ten NTD-binding antibodies tested cluster in four groups based on r values >0.5 (right). **(d)** Histogram of all deletions located in the spike protein with common NTD deletions annotated in blue. Inset: distribution of all spike substitutions, deletions, and inserts in the GISAID. **(e)** Consequences of NTD indels on ACE2 (top) and antibody (bottom) binding (n=3 biological replicates, mean ± S.D. reported in File S1).

Pearson correlation analysis of the NTD alanine and clinical variant binding data showed strong similarities between NTD binding nAbs and no correlation with REGN10987 (Fig. 3c and Supplementary Fig. 10). We clustered antibodies into four binding classes based on Pearson correlation (r) values (>0.5) between their relative binding affinities for the alanine and clinical spike mutants. The largest group consists of 4A8, 1-68, 1-87, CM17, and CM25, all of which originate from the VH1-24 multi-donor class of antibodies^47,49^. Interestingly, some antibodies derived from VH3-30 (4-18) have overlapping epitopes with VH1-24 derived nAbs but have significantly different binding modes^49^. ELISA competition assays have implicated distinct binding epitopes for 5-7 and 4-19 compared to the other antibodies in this set^44^. Our antibody classifications extend prior structure- and competition-based classifications of NTD-binding antibodies^44,49^.

Deletions and insertions in NTD loops N3 and N5 are also possible routes for immune evasion (Fig. 3d)^51,52^. A lab-evolved SARS-CoV-2 virus with an 11 amino acid insertion in N5 [_248a_KTRNKSTSRRE_248k_] evaded high titer convalescent sera^53^. To clarify the escape mechanisms, we assayed these variants against the panel of nAbs described above (Fig. 3e). Deletions within the N3 loop (Phe140-Leu141, Tyr144/145, and His146) disrupted binding to most NTD nAbs. A deletion at Phe140, located at the base of the N3 loop, also caused a moderate loss in binding for most of the NTD-binding antibodies. Similarly, a deletion at the base of the N5 loop (Ala243-Leu244) and an 11 amino acid insertion within the N5 loop resulted in reduced binding for most of the nAbs (Fig. 3e). A deletion at Ile210 had negligible effects on antibody binding. Together, these results suggest that indels within or adjacent to the N3 and N5 loops effectively abrogate nAb binding, likely due to the deletion of critical residues or spatial reconfigurations of the NTD loops. An insertion in the N5 loop also effectively evades neutralizing antibodies. As immune escape becomes a major evolutionary factor in an increasingly vaccinated world, NTD insertions may also become more common.

Next, we examined RBD mutations, eleven of which were previously reported to escape therapeutic antibodies from the REGN-COV2 cocktail (REGN10933 and REGN10987) (Supplementary Fig. 11)^35^. Expression of the full spike trimers and their escape from RBD-binding nAbs correlated well with changes in expression for the RBD alone (Supplementary Fig. 11a-e). However, K417N strongly reduced ACE2 binding in the context of the full spike relative to the isolated RBD. K417 is sterically occluded when the RBD is in the “down” position, providing a possible explanation for a larger reduction in binding observed with spike display compared to yeast display. We also examined a quaternary S1-binding nAb (2-43) and the NTD binder 4A8. As expected, 4A8 binding was not affected by RBD mutations. Three variants (L455A, F486K, and E406W) that reduced REGN10933 binding also reduced 2-43 binding (Supplementary Fig. 11b). These findings underscore the importance of characterizing nAb escape mutations within the context of the fully glycosylated spike trimer.

### Characterization of spikes from emerging variants of concern

We quantified spike expression, ACE2 affinity, and immune evasion in VOCs (Fig. 4). Four lineages, first detected in the United Kingdom (B.1.1.7), South Africa (B.1.351), Brazil (B.1.1.248), and California (B.1.427/B.1.429), harbor eight, nine, ten, and three spike mutations, respectively. Lineage B.1.1.7 increased spike expression 39% relative to the wild type (6P-D614G) spike while B.1.351, B.1.1.248, and B.1.427/B.1.429 showed 32%, 47%, and 62% reduction in spike expression, respectively (Fig. 4a). All VOCs also harbor the D614G substitution, which significantly boosts spike expression (Fig. 1d). K417N (B.1.351) and K417T (B.1.1.248) further positively compensate spike expression in some variants.

**Figure 4:**
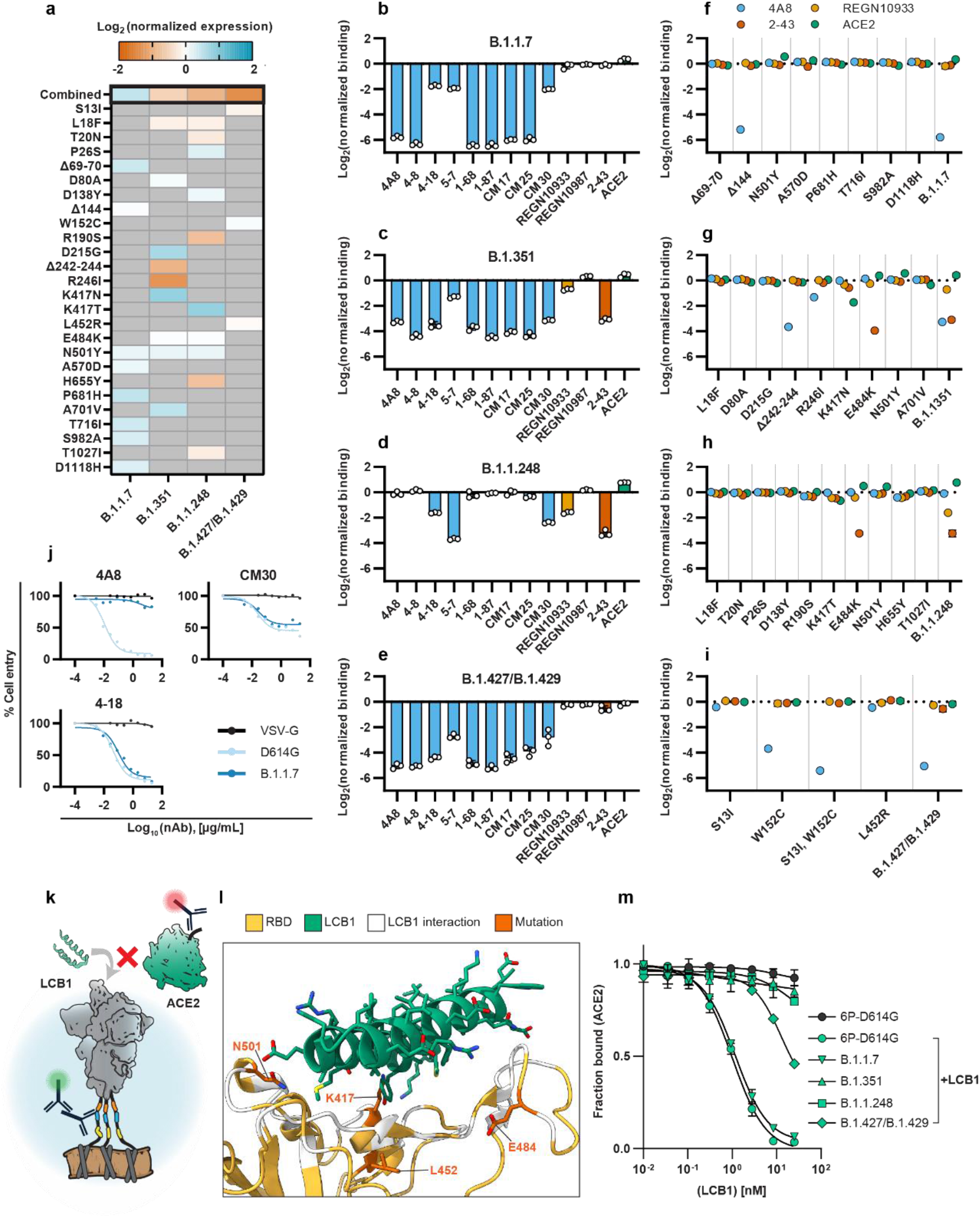
Variants of concern evade NTD-directed nAbs and ACE2 decoys. **(a)** Normalized spike expression for four variants of concern and their mutations, measured by flow cytometry. Red = decreased expression, blue = increased expression relative to spike (6P-D614G). Gray color indicates the absence of a mutation in a lineage (Mean ± S.D. of at least twelve biological replicates reported in File S1 **(b)** Relative antibody and ACE2 binding to B.1.1.7, **(c)** B.1.351, **(d)** B.1.1.248, and **(e)** B.1.427/B.1.429 as compared to spike (6P-D614G). NTD binders = blue, RBD binders = yellow, S1 binders = red, ACE2 = green. Mean ± S.D. of three biological replicates **(f-i)** Relative antibody binding measured for each variant’s mutations (n=3 biological replicates, mean ± S.D. reported in File S1) **(j)** Pseudovirus neutralization curves comparing D614G (light blue) and B.1.1.7 (dark blue) variants using three neutralizing antibodies (4A8, CM30, and 4-18). VSV-G (black) included as an infection control (also see Supplementary Fig. 13 and Methods). **(k)** Cartoon schematic of LCB1 micropeptide and ACE2 competition for RBD binding. **(l)** Co-structure (PDB: 7JZL^17^) of LCB1 (green) binding the RBD (yellow) with contacting residues (white) and four notable VOC mutations (orange) highlighted. **(m)** Competition assay with fixed ACE2 and titrated amounts of LCB1 using spike displayed VOCs and measured by flow cytometry (error bars: S.D. of three biological replicates).

We next determine the combined effect of the four VOCs on ACE2 binding and nAb escape (Figs. 4b-e). Lineages B.1.1.7, B.1.351, and B.1.1.248 bound ACE2 with a greater affinity than WT (6P-D614G), with B.1.1.248 showing the greatest increase (1.7-fold over 6P-D614G). Most NTD supersite-targeting nAbs had ∼10-fold reduced binding to the B.1.1.7, B.1.351, and B.1.427/B.1.429 lineages. Biolayer interferometry using cleaved spikes confirmed the loss of binding across most NTD-binding nAbs (Supplementary Fig. 12). However, 4-18 and CM30 both retained strong binding to B.1.1.7. Pseudovirus neutralization assays confirmed that 4-18 and CM30 can also prevent cell entry in both the reference and B.1.1.7 strains (Fig. 4j and Supplementary Fig. 13). Additionally, B.1.351 and B.1.1.248, which have three nearly identical RBD mutations showed reduced REGN10933 and 2-43 binding. We systematically screened each mutation for all VOCs using 4A8, REGN10933, 2-43, and ACE2 (Figs. 4f-i). Mutations in the N3 and N5 loops of the NTD (Δ144/145, W152C, Δ242-244, and R246I) reduced 4A8 binding. While other NTD mutations (i.e., L18F) had no detectable effect on 4A8’s binding, they evade separate classes of NTD neutralizers, such as 5-7 and 4-19. These nAbs bind predominately outside the common N3 & N5 loop supersites.

The reduction in REGN10933 binding for B.1.351 and B.1.1.248 results from the combined K417N/T and E484K mutations. Similarly, the net increase in ACE2 binding is due to a combination of E484K and N501Y offsetting the reduced ACE2 binding elicited by the K417N/T. The E484K mutation alone was sufficient to reduce 2-43 binding ∼10-fold in both strains. Mutations that reduced nAb binding (Δ242-244 and R246I) were also the most detrimental for spike expression. These results suggest that spike-destabilizing mutations that are beneficial for viral fitness (immune evasion and ACE2 binding) are compensated by other stabilizing mutations. Similar mutation patterns have been described for other viral evolution and escape pathways^54–56^.

ACE2 mimics are promising SARS-CoV-2 inhibitors but many were designed against the viral clades circulating in 2019. Their efficacy has not been assessed with modern VOCs^17,19^. We tested how these VOCs bind LCB1, an engineered alpha-helical peptide that competes with ACE2 for binding to the RBD^17^. LCB1 has a sub-nM affinity for the RBD *in vitro* and neutralizes live viruses with an IC50 of 24 pM^17^. We developed a competition assay where cells are co-incubated with increasing concentrations of LCB1 and a fixed ACE2 concentration. Changes in ACE2 binding were then measured by flow cytometry (Fig. 4k). Variant B.1.1.7 showed the same sigmoidal curve as WT (6P-D614G) spike, indicating potent inhibition of ACE2 binding above > 1 nM LCB1 (Fig. 4m). B.1.351 and B.1.1.248 showed complete escape; ACE2 binding remained unaffected even at >100 nM [LCB1]. The B.1.427/B.1.429 variant showed moderate escape. The K417N/T, L452R, E484K, and N501Y point mutations are all in the vicinity of the LCB1-ACE2 interface and are likely to reduce LCB1 binding affinity (Fig. 4l). Future peptide-based inhibitor therapies will require a cocktail of inhibitors to evade VOCs. Alternatively, these peptides must be continually optimized to improve efficacy against emerging viral variants^20^.

## Discussion

We developed spike display to accelerate vaccine design and to rapidly assess the effects of mutations in emerging virus strains. Pre-fusion stabilized spike is the primary antigen delivered via immunization. Pre-fusion stabilization requires laborious biochemical assays following structure-guided selection of promising stabilizing mutations^12,38,40,57^. Spike display shortens this workflow by rapidly assessing the expression of individual point mutants and their combinatorial derivatives (Fig. 1d). Spikes can be cleaved from cell surfaces for structural and biophysical characterization. We anticipate that spike display can be adapted for diverse coronavirus-family spikes and other viral antigens (Fig. 1e)^32,58,59^.

Spike display complements structure-function studies for high-resolution epitope mapping. We focused on the NTD because it is under heavy mutational pressure in circulating SARS-CoV-2 variants and elicits neutralizing antibodies in convalescent patients^43,44,47,50,60^. The mechanisms of NTD-mediated viral inhibition remain unexplored, but these antibodies do not block ACE2 binding. Our NTD scan uncovered that most of the NTD-binding nAbs recognize a public epitope comprised of the N3/N5 loops. These loops may be required for additional conformations that occur after ACE2 recognition. Although circulating variants escape first generation NTD-binding nAbs, these epitopes are an important target for second generation antibody therapeutics. We conclude that the NTD is under strong evolutionary pressure and that existing NTD-targeting nAbs must be paired with nAbs that target other epitopes to avoid therapeutic evasion.

Using spike display, we screened variants B.1.1.7 (UK), B.1.351 (S. Africa), B.1.1.248 (Brazil), and B.1.427/B.1.429 (CA), and their individual mutations (Fig. 4). The B.1.1.7, B.1.351, and B.1.1.248 lineages share a common N501Y RBD mutation that likely confers greater transmissibility^27,62,63^. The B.1.427/B.1.429 variant also exhibits improved transmissibility relative to the parental strain^30^. Compensatory mutations offset the de-stabilizing effects of NTD and RBD mutations that partially escape NTD- and RBD-binding nAbs. Variants B.1.1.7, B.1.351, and B.1.427/B.1.429 all escaped NTD-binding nAbs. However, many NTD nAbs are promising therapeutics for B.1.1.248, which does not have any consequential mutations near the NTD supersite. The RBD-binding REGENERON antibody cocktail recognized all variants, with B.1.1.248 showing a modest reduction in binding due to the K417T and H655Y substitutions. B.1.1.7 and B.1.351 also completely evaded a promising micro-peptide ACE2 mimic. Future ACE2 mimetic therapies will need to be re-formulated to stay ahead of current and emerging variants of concern.

Spike display accelerates the characterization of antibody-spike interactions and the consequences of non-synonymous spike mutations on these interactions. The entire process, from Golden Gate assembly to flow-based characterization can be completed in < 5 days. Downstream biophysical studies can use the cell-cleaved spikes without laborious recombinant protein purification. A second-generation spike display platform will increase throughput by integrating spike variants in a chromosomal locus and sorting pooled libraries for phenotypic differences^64–66^. Increasing throughput will allow interrogation of all circulating non-synonymous spike mutations and rapid protein engineering for pre-fusion stabilization. Deep mutational scanning of all possible amino acid substitutions will be broadly useful for characterizing the spike mutational landscape as well the development of pan-coronavirus vaccine antigens.

## Acknowledgments

We thank Drs. Ching-Lin Hseih, Jason S. McLellan, Jason Lavinder, and Gregory C. Ippolito for generously sharing ACE2 and antibody expression constructs, and purified antibodies. We are grateful to members of the Finkelstein & Ippolito laboratories for carefully reading the manuscript. This work was supported by the Welch Foundation (F-1808 to I.J.F.; F-1390 to K.N.D.), the Bill and Malinda Gates Foundation (INV-017592 to I.J.F.), and a Cooperative Agreement between UT Austin and ARL (W911NF-17-2-0091 to J.D.G.).

## Author Contributions

K.J., C.-W.C., J.D.G., and I.J.F. designed the research. K.J. performed the flow experiments. K.J., C.-W.C., C.I.T., A.A., Q.G., G.N., D.R.B., J.G., W.N.V., and H.-C.K. purified antibodies and other reagents. K.J., C.I.T., A.A., Q.G, J.L., H.Z., cloned spike variants. T.S.K and K.N.D. provided the pseudovirus neutralization data. K.J., C.-W.C., C.I.T., H.Z., and A.P.H. analyzed the data. K.J. and I.J.F. wrote the paper with editorial assistance from all co-authors.

## Data Availability

All relevant raw data are available from the corresponding authors upon reasonable request.

## Materials and Methods

Oligonucleotides and gene blocks were purchased from Integrated DNA Technologies (IDT). Restriction enzymes, polymerases, and ligases were purchased from New England Biolabs (NEB). Antibodies were purified as described below or purchased from the following manufacturers: Mouse anti-FLAG M2 (Sigma-Aldrich; F3165); Goat anti-Mouse IgG(H+L), Human ads-Alexa Fluor® 488 (SouthernBiotech; 1031-30); Goat anti-Human IgG Fc-Alexa Fluor® 647(SouthernBiotech; 2048-31); Goat anti-Mouse IgG H&L (IRDYe® 680RD) (abcam; ab216776); Beta Tubulin Loading Control Monoclonal Antibody (BT7R) (Thermo-Fisher; MA5-16308)

### Mammalian display vectors and spike protein mutants

Spike display plasmids incorporate the pre-fusion stabilized SARS-CoV-2 S-6P (“HexaPro”) as the reference sequence for all spike variants (Addgene #154754)^38^. This construct comprises residues 1-1208 of SARS-CoV-2 S gene (GenBank: MN908947) with prolines substituted at residues F817, A892, A899, A942, K986, V987, a protease-inactive furin cleavage site (^682^GSAS^685^), and the globally dominant D614G mutation^23^. This construct (6P-D614G) was optimized for mammalian cell surface display and high throughput Golden Gate cloning as described below.

For mammalian surface display, spike was cloned into a pcDNA5-based vector (Addgene #113547) with the following additional N-terminal Ig-Kappa leader and C-terminal epitopes ordered as two separate gBlocks (IDT): a 3X-FLAG, a Strep Tag II epitope, an HRV 3C protease site, and a PDGFR-β transmembrane domain. All three parts (N-terminal leader, spike, and C-terminal additions) were assembled with the pcDNA5 backbone using Hi-Fi DNA assembly (NEB; E2621S).

To enable high throughput Golden Gate cloning of cell surface-displayed spike variants, the spike coding region was divided into 5 parts with junctions strategically positioned at amino acids with low mutational frequencies, according to the GISAID. For each of the 5 parts, an entry vector was constructed by cloning in a superfolder GFP (sfGFP) bacterial expression cassette with flanking AarI cut sites and unique 4 nt overhangs matching the wild-type SARS-CoV-2 sequence of each part junction, using PCR and Hi-Fi DNA assembly (See Supplementary Fig. 2). A part 1-5 entry vector was made with the entire SARS-CoV-2 coding sequence replaced with the sfGFP cassette to enable multi-part assemblies of complex spike variants or entirely different spike proteins.

Parts 1,2,3,4, and 5 of the 6P-D614G coding sequence were PCR amplified with flanking AarI cut sites, the matching 4 nt overhangs for spike assembly (See Supplementary Fig. 2), Esp3I cut sites, and 4 nt overhangs for YTK001 assembly. The Esp3I (NEB; R0734) and adjacent 4 nt overhangs were used to incorporate, via Golden Gate cloning, each part (1-5) coding sequences into the YTK001 entry vector from the yeast toolkit^68^.

### Automated spike variant cloning pipeline

Golden Gate constructs were assembled using a high-throughput automated pipeline that includes an Echo 525 Acoustic Liquid Handler, a Tecan Fluent, and a QPix 420 Colony Picker. Golden Gate compatible parts were arranged in a 384-well Echo Source Plate (PP-0200) and transferred to 96-well PCR destination plates using the Echo 525. Each well of the 96-well destination plate received the following Golden Gate reaction mixture: 0.5 µL of T7 DNA Ligase (NEB; M0318S), 0.5 µL of AarI (Thermo Fisher; ER1582), 0.2 µL AarI Oligo (Thermo Fisher), 1 µL T4 DNA Ligase Buffer (NEB; B0202A), 1 µL of the insert and plasmid DNA, and nuclease-free water to bring the final volume to 10 µL per reaction.

Reaction mixtures were incubated on a thermocycler according to the following conditions: 25 cycles of digestion and ligation (37 °C for 1 min, and 16 °C for 2 mins), followed by a final digestion (37 °C for 30 mins), and a heat inactivation step (80 °C for 20 mins). To improve assembly efficiencies for assemblies with 4+ parts increase the cycled digestion and ligation steps to 3 min and 5 min, respectively.

96-well PCR plates with 50 µL of DH10B ‘Mix & Go Competent Cells’ (prepared using Zymo T3001) in each well were prepared beforehand so that high-throughput transfers could be done using multichannel pipettes or the Tecan Fluent. To transform the cells, we transferred 4 µL from each unique reaction mixture to respective wells containing 50 µL of Mix & Go competent cells. Wells were mixed by pipetting and the cells were incubated at 4 °C for 10 minutes. The DNA-cell mixtures were then transferred to an Axygen deep well grow block (P-2ML-SQ-C-S) with 150 µL of superior broth (AthenaES; 0105) in each well and incubated for 1 hour at 37 °C with shaking at 950 rpm on a plate shaker.

Outgrown cells were plated dropwise on Nunc OmniTrays (5 μL per plate) (Thermo Fisher; 140156), containing LB-agar + carbenicillin at 100 µg/mL. Each plate can fit 96, 5 µL drops. Plates were kept at room temperature until the drops dried and were then transferred to a 37 °C incubator for growth overnight (12-16 hours).

Colonies were screened the following day and picked using the QPix 420, selecting only white colonies and avoiding green fluorescent colonies, which still contain the sfGFP cassette and not the desired spike sequence. Colonies were picked into 1 mL of SB media with antibiotic in Axygen deep well grow blocks and grown overnight at 37 °C while shaking. Once grown, liquid cultures were pelleted at 3000 g for 10 minutes and miniprepped using the Tecan Fluent robotic liquid handler with Promega Wizard SV 96 Plasmid DNA Purification Kit (Promega; A2250).

### Expression and purification of neutralizing anti-spike antibodies

Previously published VH and VL sequences were purchased as gBlocks (IDT) to create full-length antibody IgGs^43–45^. VHs and VLs were cloned into custom Golden Gate compatible pcDNA3.4 vectors for IgG1 expression.

Expi293 cells were cultured in Expi293 Expression Medium (Sigma-Aldrich; A1435101) and maintained in a humidified atmosphere of 8% CO_2_ and 37 °C while shaking continuously at 125 rpm. Cells were transfected with a 1:3 molar ratio of heavy chain and light chain expression vectors using the Expi293 Transfection kit (Sigma-Aldrich; L3287) according to the manufacturer’s instructions. Five days after transfection, the protein-containing supernatant was collected by two centrifugation steps. Cells and supernatant were first separated by spinning cultures at 300 g for 5 min at 4 °C. Cell debris and supernatant were separated by spinning at 3,000 g for 25 min at 4 °C. To purify human IgGs, Protein G magnetic beads (Promega; G7471) were washed with PBS buffer and then added to the supernatant in a 1:10 volumetric ratio. After incubating supernatant and bead mixtures for 1 hr with gyration at room temperature, bead-bound antibodies were pelleted on a magnetic peg stand for washing and final elution with 100 mM glycine-HCl (pH 2.5). Residual beads were clarified by running the elute through a 0.22 μm syringe filter, and then neutralized with 2 M Tris buffer (pH 7.5). Purified antibodies were kept at 4 °C for short-term storage, and frozen at −20 °C for long-term storage.

### Expression and purification of chimeric ACE2-Fc

Human ACE2-Fc was recombinantly expressed in Expi293 cells with minor modifications to previously published protocols^12^. Simply, the ACE2-Fc expression vector was transfected into Expi293T cells according to the manufacturer’s instructions (Sigma-Aldrich; L3287). Five days after transfection the supernatant was collected by first spinning cultures at 300 g for 5 min at 4°C. Cell debris and supernatant were further separated by spinning at 10,000 g for 20 min at 4°C. Upon resuspending in PBS, ACE2-Fc was purified over Protein A Agarose (Thermo Fisher; 15918014). Once the Protein A Agarose was equilibrated in PBS buffer, the respective supernatant was applied 3X and washed with 10 bed volumes of PBS buffer. The protein was eluted with 100mM glycine-HCL (pH 2.4) into 0.1X volume Tris buffer (pH 8.5) and 100 mM NaCl. Purified ACE2-Fc was stored at 4 °C for short-term storage, and frozen at −20 °C for long-term storage.

### Expression and purification of SARS-CoV-2 spike proteins

Plasmids were transfected into Expi293 cells and expressed as described previously^38^. Variants were purified from 40mL cell culture. The supernatant was filtered by 0.22 μm filter and then run over StrepTactin Superflow column (IBA 2-1206-025). Spikes were further purified by Superose 6 increase 10/300 (GE healthcare) size-exclusion column in a buffer containing 2mM Tris pH 8.0, 200mM NaCl, and 0.02% NaN_3_. Purified samples were stored at 4 °C for short-term storage, and frozen at −20 °C for long-term storage.

### HEK293T culturing and transfection

HEK293T cells were cultured in DMEM (Gibco; 11995065) containing phenol red, 4 mM L-glutamine, 110 mg L-1 sodium pyruvate, 4.5 g L-1 D-glucose, and supplemented with 10% FBS (Gibco; 26140079) and 2% Pen/Strep (Thermo Fisher; 15070063). Cells lines were authenticated and tested for mycoplasma contamination before use via the Mycoplasma Detection Kit (SouthernBiotech; 13100-01). Cells were maintained in a humidified atmosphere of 5% CO_2_ and 37 °C and were passaged every 2-3 days into 10 cm polystyrene coated plates (Eppendorf; EP0030700112-300EA) upon reaching high density. Approximately 24 hrs before transfection, cells were seeded into 6-well or 12-well polystyrene coated plates (Eppendorf; EP0030720130, EP0030721012) at a density of 0.3 x 10^6^ cells mL^−1^ or 0.1 x 10^6^ cells mL^−1^, respectively. Upon reaching 60-80% confluence, cells were transfected with expression plasmids using Lipofectamine 3000 (Thermo Fisher; L3000015) following the manufacturer’s instructions and 3 μL of lipofectamine per μg of plasmid DNA. Cells were assayed or collected 48 hrs post-transfection.

### Flow cytometry and data analysis

HEK293T cells with surface-displayed spike were collected 48 hrs post-transfection by first washing once with PBS and then resuspending in PBS by gentle pipetting. Cell density was determined using a cell counter (Logos Biosystems; L40002). Cells were then spun down at 200 g for 1 min. After decanting the supernatant cells were resuspended in chilled PBS-BSA (1% BSA, 1X PBS, 2 mM EDTA, pH 7.4) to a density of ∼3 x 10^6^ cells mL^−1^.

Flow cytometry assays were prepared using an Axygen Deep well grow block (P-2ML-SQ-C-S). Each well contained a predetermined concentration of primary antibody or chimeric cell receptor (ACE2-Fc) diluted in PBS-BSA and 50 μL (1.5 x 10^5^ cells). Mixtures were incubated at room temperature and shaken at 950 rpm for 1 hr. Cells were then pelleted by spinning the plate for 2 min at 500 g in a swinging bucket rotor. Cells were washed twice by decanting the supernatant and adding 500 μL of PBS-BSA to each well. 500 μL of a secondary antibody solution (5 μM Alexa Fluor® 488 anti-mouse (SouthernBiotech; 1031-30) and 10 μM Alexa Fluor® 647 anti-human (SouthernBiotech; 2048-31) in PBS-BSA) was added to each well. The plate was incubated in the dark, at 4 °C while shaking at 950 g for 25 min. Wells were washed again, twice with PBS-BSA, and then resuspended in 300 uL of PBS-BSA before running on the SA3800 Spectral Analyzer (SONY).

Authenticated HEK293T cells were used to establish forward scatter-area (FSC-A) and side scatter-area (SSC-A) gating. Singlet discrimination was then established with forward scatter-height (FSC-H) vs forward scatter-area (FSC-A) and side scatter-height (SSC-H) vs side scatter-area (SSC-A) gates. A minimum of 10,000 singlet events were acquired for each assayed sample. These singlet HEK293T cells were further analyzed in two fluorescent channels, Alexa Fluor 488 and Alexa Fluor 647, using manufacturer-recommended excitation and detection settings. Spectral unmixing was applied to all data to reduce the effect of spectral spillover and autofluorescence on downstream calculations.

Median height (H) measurements for the AF-488 and AF-647 channels were recorded for each sample. Anti-FLAG (AF-488 channel) signal was used to measure spike expression. Spike variant (x) expression relative to WT (6P-D614G) was calculated using the following equation:

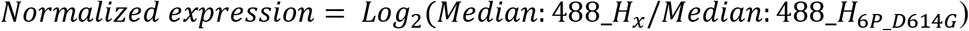

Anti-FLAG signal was also used as an internal normalization control to correct for changes in transfection efficiency and spike expression when measuring antibody or ACE2 binding. Normalized binding measurements for spike variants (x) expression relative to WT (6P-D614G) was calculated using the following equation:

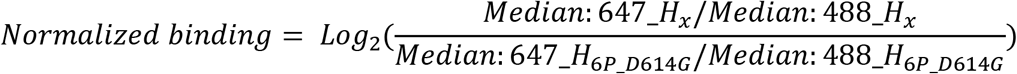

All flow cytometry data were analyzed using FlowJo v9.

### LCB1 cloning and purification

LCB1 was synthesized and cloned into E.coli using a pET expression vector via Hi-Fi DNA assembly (NEB; E2621S) with a gBlock (ITD) encoding an N-terminal 8x His tag followed by a TEV cleavage site and the LCB1 gene sequence^17^. The 8xHis-LCB1 was transformed into homemade chemically competent Rosetta (DE3) cells. Protein expression was performed using LB media supplemented with Carbenicillin, induced with IPTG and grown overnight. Cells were pelleted at 4,000 g for 10 minutes and resuspended in lysis buffer containing 30 mM Tris-HCl, pH 7.5 and 300 mM NaCl with DNAse and protease inhibitor tablets. The cells were lysed by sonication while on ice. Soluble material was then clarified by centrifugation at 35,000 g for 45 minutes. The soluble fraction was added to a 5 mL HisTrap HP column (GE Healthcare; 17524802) and eluted with elution buffer containing 30 mM Tris-HCl, pH 7.5, 300 mM NaCl, and 250 mM Imidazole. The 8xHis-LCB1 was dialyzed overnight with TEV protease at 4 °C in buffer containing 30 mM Tris-HCl, pH 7.5 and 300 mM NaCl. The TEV protease and 8xHis tag was removed by running the protein on a 5 mL HisTrap HP column and collecting the flowthrough.

### LCB1 and ACE2 competition assays

Spike displaying HEK293T cells were collected and incubated with varying concentrations of LCB1 peptide and a fixed (25 nM) ACE2-Fc concentration. Anti-FLAG (Sigma-Aldrich; F3165) was also included at 1 μg/mL final concentration. Cells were incubated at room temperature in an Axygen Deep well grow block (P-2ML-SQ-C-S), shaking at 950 rpm for 1 hr. Cells were then washed with PBS-BSA and stained with secondary antibodies (5 μM Alexa Fluor® 488 anti-mouse (SouthernBiotech; 1031-30) and 10 μM Alexa Fluor® 647 anti-human (SouthernBiotech; 2048-31) in PBS-BSA), shaking at 950 rpm for 1 hr at 4°C. After two final washes with PBS-BSA, cells were resuspended in PBS-BSA and run on the SA3800 Spectral Analyzer (SONY). Data were analyzed using FlowJo v9.

### Cell immunostaining and microscopy

HEK293T cells were seeded into imaging dishes (Eppendorf; 0030740009) and transfected with spike display plasmids. After 48 hrs, cells were gently washed twice with PBS and then fixed with 4% formaldehyde for 20 min at room temperature. Cells were washed again with PBS and then blocked with PBS-BSA (1%) blocking buffer for 20 min. Primary antibodies (anti-FLAG M2; Sigma and chimeric ACE2-Fc (Genscript; Z03484) were diluted in PBS-BSA to 1 μg mL^−1^ and added to each imaging dish, incubating at room temperature for 1 hr. Cells were washed twice with PBS-BSA. Secondary antibodies (Alexa Fluor® 488 (SouthernBiotech; 1031-30); Alexa Fluor® 647(SouthernBiotech; 2048-31) were diluted in PBS-BSA to 10 μg mL^−1^ and added to each imaging dish, incubating at room temperature for 1 hr in complete darkness. Cells were washed twice with PBS-BSA. Hoechst stain (Thermo Fisher; 62249) was diluted 1:10,000 in PBS-BSA and added to each imaging dish, incubating at room temperature for 15 min. Cells were washed once with PBS-BSA and a final 1 mL of PBS-BSA was added before imaging.

All images were collected with a Nikon Ts2R-FL inverted research microscope equipped with a CF160 Plan Apochromat Lambda 60x oil immersion objective lens. Hoechst stain, Alexa Fluor® 488, and Alexa Fluor® 647 were excited by a SOLA SM II 365 light engine and filtered through C-FL Hoechst (AT460/50m), C-FL FITC (AT535/40m), and C-FL mCherry (ET630/75m) filter cubes (Chroma), respectively. Images were acquired with a pco.panda sCMOS camera controlled with NIS-Elements D software.

### SARS-COV-2 pseudovirus neutralization assay

Most of the plasmids needed for expressing the HIV virion under the CMV promoter were obtained from BEI resources, including HDM-Hgpm2, pRC-CMV-Rev1b, and HDM-tat1b (BEI catalog numbers NR-52516, NR-52519, and NR-5251, respectively)^69^. The lentiviral backbone plasmid, expressing a luciferase reporter under the CMV promoter followed by an IRES and ZsGreen, was also provided by the BEI resources (NR-52520). The envelope plasmid (HDM-IDTSpike-fixK) expresses a codon-optimized WT spike protein of SARS-COV-2 under a CMV promoter (Genbank ID: NC_045512) and was supplied by BEI resources. It was used as a template for site-directed mutagenesis to generate new expression plasmids for the D614G and B.1.1.7 variants. The plasmid expressing the VSV-G (vesicular stomatitis virus glycoprotein) was purchased from Cell Biolabs (pCMV-VSV-G, Part No. RV-110). The lentiviral plasmid used to generate the HEK-293T stable cell lines expressing the human ACE2 gene (GenBank ID NM_021804) under an EF1a promoter was obtained from the BEI resources as NR52516^69^.

The HIV particles pseudotyped with SARS-CoV-2 spike variants D614G or B.1.1.7 were generated in HEK293T cells, following previously published protocols^69^. Cells were transiently co-transfected with plasmids for (1) HIV virion formation proteins (HDM-Hgpm2, pRC-CMV-Rev1b, and HDM-tat1b; (2) one of the envelope proteins (2019-nCoV Spike-D614G mutant, B.1.1.7 variant or VSV-G) and (3) the lentiviral backbone expressing luciferase reporter (pHAGE-CMV-Luc2-IRES-ZsGreen-W). Media was exchanged for new media 24 hours post-transfection. Media containing the pseudovirus particles were collected, filtered, and fractionated 72 hours post-transfection. Fractions were stored at −80 °C until later usage. Lentivirus titer was estimated for each virus particle using RT-PCR, following the manufacturer’s protocol (qPCR Lentivirus Titer Kit, Applied Biological Material Catalogue LV-900). Around 106 infectious units of the virus were incubated with each dose of the tested antibody in full media (100 µL) for 1 hour at room temperature. The mixture was then added to HEK-293T target cells, stably expressing human ACE2 in 96-well white plates with a clear bottom. The % cell confluency in each well was estimated after 60 hours of incubation using the lncuCyte® ZOOM equipment with a ×10 objective. Cells were then treated with Bright-Glo Luciferase Assay System (Promega; E2610) to detect luciferase signal (relative luciferase units or RLU) following the manufacturer protocol. The ratio of recorded relative luciferase units in the antibody’s presence to the readout in the absence of the antibody was employed to estimate the percentage of virus entry at each dose. Half-maximal inhibitory concentrations (IC50) were calculated using a 3-parameter logistic regression equation (GraphPad Prism v9.0). Experiments were performed in duplicate using different preparations of virus.

### Western blots

Equal cell numbers were collected 48 hrs post-transfection for each sample. Proteins were extracted using RIPA buffer with protease inhibitors (cOmplete™, EDTA-free Protease Inhibitor Cocktail; Millipore Sigma). Sample input was normalized using a Bradford protein assay that was calibrated with BSA standards. For each sample, 10 μg of total protein was run on a 4–15% SDS-PAGE gel (Mini-PROTEAN TGX Precast Protein Gel; Bio-Rad). Proteins were then transferred onto a PVDF membrane using a Trans-Blot® SD Semi-Dry Transfer Cell (Bio-Rad) per the manufacturer’s instructions. Membranes were blocked in LICOR Odyssey Blocking Buffer (Neta Scientific) incubated with the following primary antibodies overnight: mouse anti-FLAG M2 antibody (1:10,000; Sigma-Aldrich; F3165) for spike protein or RBD detection and mouse anti-beta tubulin antibody (1:5000; Thermo Fisher; MA5-16308) as a loading control. The membrane was washed three times with PBST (1X PBS, 0.1% Tween 20) for 10 min and then incubated with the following secondary antibody for 2 hrs: goat anti-mouse IRDYe® 680 conjugated antibody (1:10,000; abcam; ab216776). The membrane was washed again with PBST three times for 10 min and imaged on the Odyssey Li-Cor.

### Spike display cleavage and spike purification

HEK293T cells were transfected with plasmids encoding spike display variants and collected 48 hrs post-transfection by washing once with PBS and resuspending 3-4 x 10^6^ cells in a 1.5 mL Eppendorf tube in 1 mL of 3C cleavage buffer (150 mM NaCl, 50 mM Tris-HCl pH 8.0). Five units of 3C protease (Thermo Fisher; 88946) were added to each tube and tubes were placed on a shaker at room temperature and 900 rpm for 1 hr to separate spike trimers from the cell surface. Supernatant containing the spike protein was collected by spinning tubes at 16,000 g for 1 min and transferring ∼1 mL of supernatant to a fresh tube. Supernatants were kept on ice until analysis. For EM imaging, supernatants were purified through a 0.5 mL StrepTactin resin (IBA) column. After spin concentrating the elution fractions to 1 mL, spikes were further purified by size-exclusion chromatography using a Superose 6 Increase 10/300 column (GE Healthcare).

### Negative stain EM data collection and processing

Purified SARS-CoV-2 spike variants were diluted to a concentration of 40 μg mL^−1^ in 50 mM Tris-HCL pH 8.0 and 150 mM NaCl. CF400-CU grids (Electron Microscopy Science) were cleaned in a Gatan Solarus 950 plasma cleaner for 45 seconds. Spikes were deposited on the girds and stained with Nano-W^®^ (Methylamine Tungstate). Grids were then imaged on a Talos F200C TEM microscope equipped with a Ceta 16M detector (Thermo Fisher). 106 micrographs were imaged from a single grid at a magnification of 92000X, corresponding to a calibrated pixel size of 1.63 Å/pix. Motion correction, CTF estimation and particle picking were performed in CisTEM^70^. Particle stacks were then imported into cryoSPARC v3.1.0 for 2D classification, *ab initio* 3D reconstruction, heterogeneous 3D refinement, and homogeneous 3D refinement^71^.

### Biolayer Interferometry

After Cleavage with 3C protease, supernatants containing spikes variants were diluted 2-fold with BLI buffer composed of 10 mM HEPES pH 7.5, 150 mM NaCl, 3 mM EDTA, 0.05% v/v Surfactant P20, 1 mg/mL bovine serum albumin. Analytes were also serial diluted with BLI buffer. Anti-mouse Fc capture (AMC) biosensor (FortéBio) were hydrated with BLI buffer for 10 mins in an Octet RED96e (FortéBio). Then, anti-FLAG IgG which is produced by mouse was immobilized to the AMC sensor tip. The assay went through the following steps: 1) baseline: 60 s with BLI buffer; 2) IgG immobilizing: 360 s with anti-FLAG IgG; 3) spikes loading: 360 s with diluted supernatants; 4) baseline: 300 s with BLI buffer; 5) association: 600 s with serial diluted analytes (antibodies or ACE2); 6) dissociation: 600 s with BLI buffer. The data were reference-subtracted and analyzed by Octet Data Analysis software v11.1 using steady-state analysis. All structures (7DDN^67^, 7C2L^43^, and 7JZL^17^) were downloaded from the RCSB-PDB as PDB files and imported into ChimeraX 1.1 for visualization and image creation.

### Computational analysis of GISAID sequence data

To identify and obtain SARS-CoV-2 spike protein variants of interest, we performed pairwise sequence alignment of all spike proteins in the GISAID EpiCoV database and the SARS-CoV-2 reference sequence with the genbank number BCN86353.1^48^. We downloaded all published spike protein sequences from the GISAID (accessed on 2/24/2021) as a FASTA file. We then used the Biopython Bio.SeqIO package (https://biopython.org/wiki/SeqIO) to extract the spike protein amino acid sequence and related species information from each entry in the FASTA file. Preliminary filtering was performed to remove non-human sequences, sequences with more than 800 unknown amino acid sequences (“X”), and sequences with <1250 or >1280 amino acids. 484,744 out of the 606,304 sequences passed these filters. GB number BCN86353.1^48^.

We then used the python parasail package (https://github.com/jeffdaily/parasail-python) to perform a semi-global alignment of each sequence to the reference sequence^72^. Parameters were set to no penalties for gaps at the beginning and end of both sequences using a BLOSUM80 substitution scoring matrix, a gap opening penalty of 5 within the sequence, and a gap extension penalty of 2 within the sequence. A BLOSUM80 scoring matrix was used because higher number blosum matrices (blosum80 as opposed to blossom32) compare closely related sequences. The highest scoring sequence alignment was kept and the aligned sequence was compared position-wise to the reference sequence to identify insertions, deletions, and substitutions. Frequencies of each unique substitution, insertion, and deletion were obtained with the following equation:

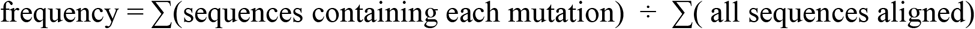

Data analysis was performed with the R tidyverse package and ggplot2^73^.

### Statistical analyses

The means ± S.D. were calculated and reported for all appropriate data. NTD-targeting nAbs were group based on Pearson correlation r values >0.5.

