## Supplementary Figures for "Rapid characterization of spike variants via mammalian cell surface display"

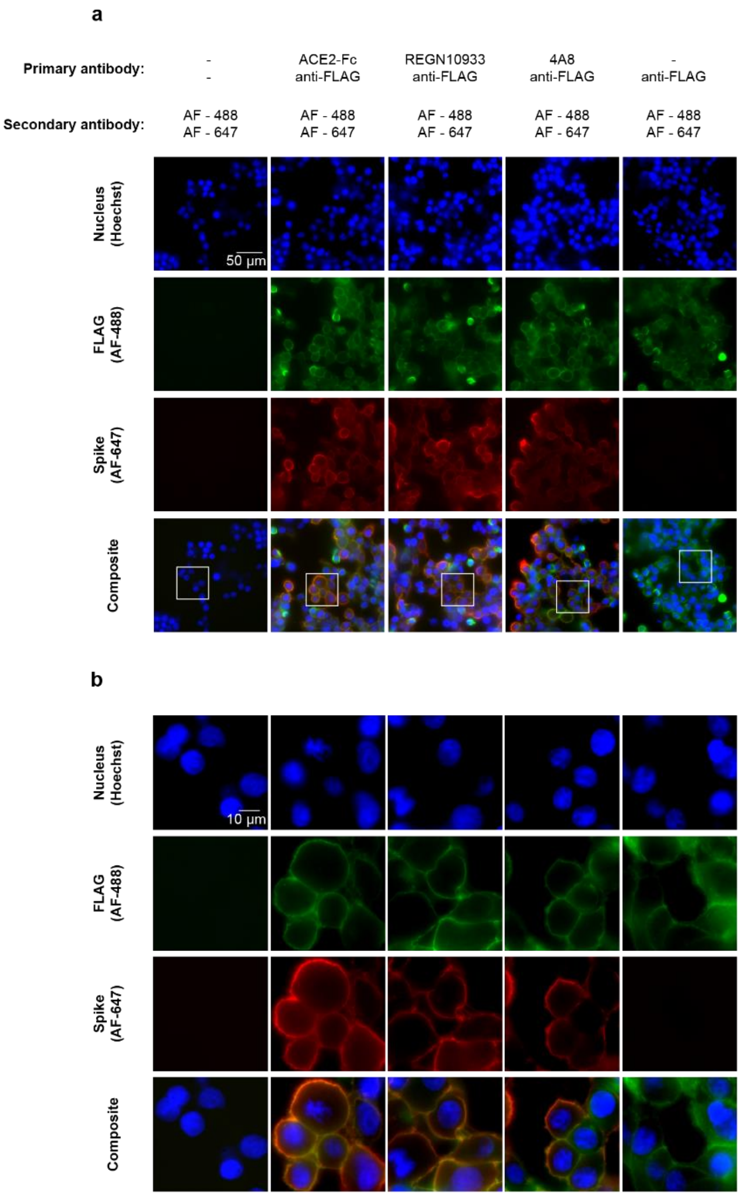


**Supplementary Fig. 1. Immunostaining shows spike on cell surfaces.**
Fluorescent immunostaining of HEK293T cells transiently expressing SARS-CoV-2 spike (6P-D614G). Cells were immunostained 48 hrs post transfection with the spike display construct. **(a)** Full field view of HEK293Ts at 60X magnification. Top row indicates how cells were immunostained; the first column is a negative control. Last column shows no AF-647 cross reactivity with anti-FLAG antibody. **(b)** Higher magnification images from the white rectangular frames in (a). AF = Alexa Fluor

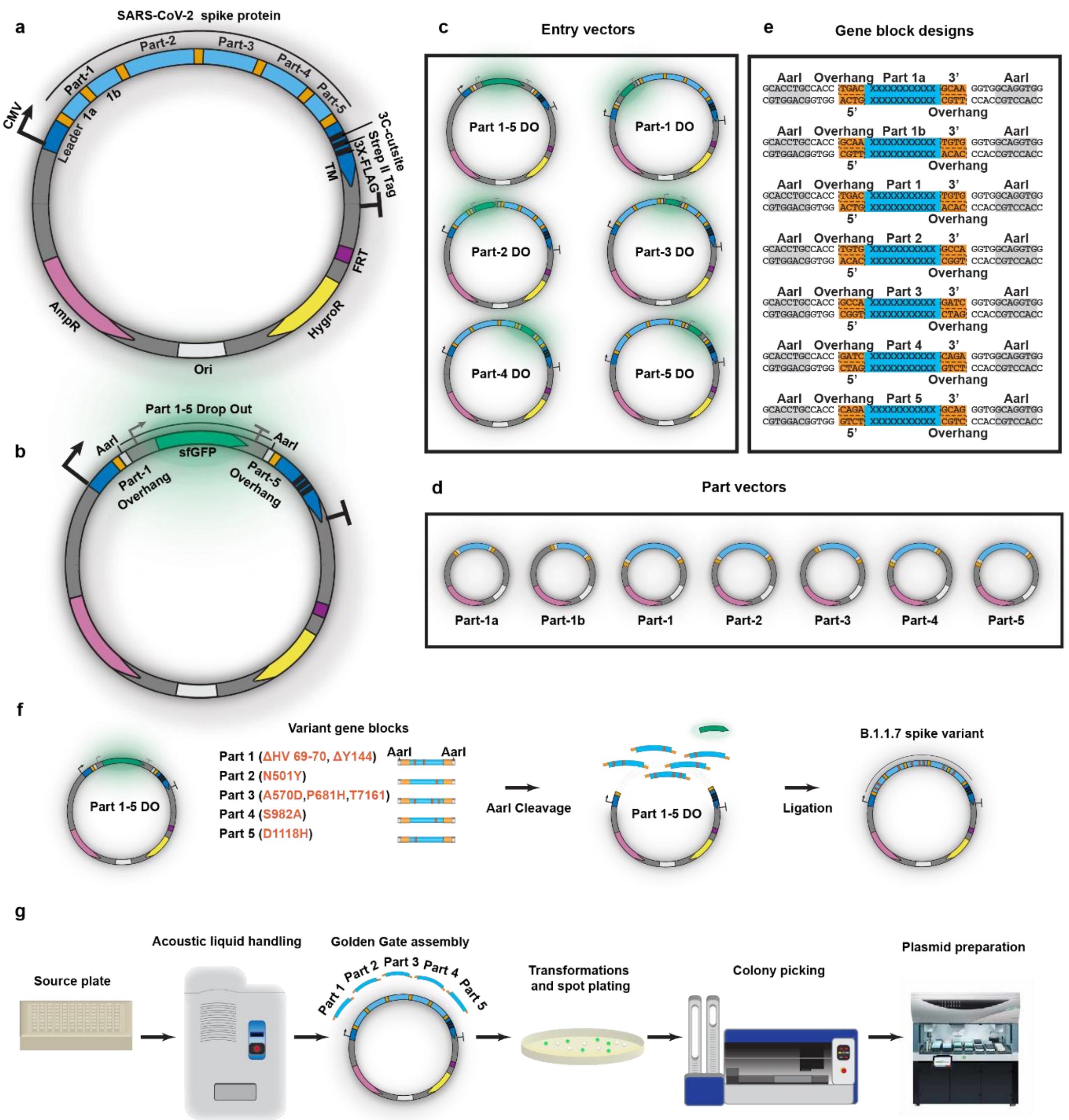

a
SARS-CoV-2 spike protein
Part-1
Part-2
Part-3
Part-4
Part-5
CMV
Leader
1a
1b
3C-cutsite
Strep II Tag
3X-FLAG
TM
FRT
HygroR
AmpR
Ori
b
Part 1-5 Drop Out
AarI
sfGFP
AarI
Part-1 Overhang
Part-5 Overhang
c
Entry vectors
Part 1-5 DO
Part-1 DO
Part-2 DO
Part-3 DO
Part-4 DO
Part-5 DO
d
Part vectors
Part-1a
Part-1b
Part-1
Part-2
Part-3
Part-4
Part-5
e
Gene block designs
AarI Overhang Part 1a 3' AarI
5' Overhang
AarI Overhang Part 1b 3' AarI
5' Overhang
AarI Overhang Part 1 3' AarI
5' Overhang
AarI Overhang Part 2 3' AarI
5' Overhang
AarI Overhang Part 3 3' AarI
5' Overhang
AarI Overhang Part 4 3' AarI
5' Overhang
AarI Overhang Part 5 3' AarI
5' Overhang
f
Variant gene blocks
Part 1-5 DO
Part 1 (ΔHV 69-70, ΔY144)
Part 2 (N501Y)
Part 3 (A570D,P681H,T716I)
Part 4 (S982A)
Part 5 (D1118H)
AarI
AarI
AarI Cleavage
Part 1-5 DO
Ligation
B.1.1.7 spike variant
g
Source plate
Acoustic liquid handling
Golden Gate assembly
Part 1
Part 2
Part 3
Part 4
Part 5
Transformations and spot plating
Colony picking
Plasmid preparation

**Supplementary Fig. 2. Automated Golden Gate assembly of spike variants. (a)** Schematic of the spike display mammalian vector with an Ig Kappa leader sequence, 3X-FLAG epitope, Strep-II affinity tag, 3C protease cleavage site, and PDGFR-β transmembrane domain (TM) driven by a CMV promoter. The spike coding region is divided into five parts (light blue) separated by unique four nt junctions (orange). Part-1 is further subdivided into Part-1a and Part-1b for high throughput cloning of NTD variants. Ampicillin resistance (AmpR), Hygromycin resistance (HygroR), and a FRT recombination site are included in the plasmid backbone. **(b)** Spike display drop-out vectors include a superfolder Green Fluorescent Protein (sfGFP; green) cassette flanked by AarI cut sites[1]. **(c)** Entry vectors enable high-throughput integration of gene blocks to rapidly produce spike variants. The vectors include a sfGFP drop-out insert flanked by parts of the wild type spike. The sfGFP cassette is flanked by unique upstream and downstream AarI-generated overhangs. This makes the gene blocks harboring specific point mutations interchangeable, allowing rapid assembly of multi-site spike mutants. **(d)** Each part vector contains the WT spike sequence corresponding to the respective part type. The unique overhang pairs for spike part within the part vectors are complementary to the corresponding entry vectors (i.e., entry Vector part-1 DO is compatible part vector part-1). **(e)** Gene block overhangs for each of the entry vectors. **(f)** Assembly of the B.1.1.7 spike variant of concern is demonstrated as an illustrative example. **(g)** Cloning workflow: An acoustic liquid handler (Echo) transfers selected parts from source plate to destination plate to assemble the Golden Gate reaction. The reaction is plated on a LB-agar + Amp plate, and white colonies are selected via the QPix Colony Picker. The selected colonies are miniprepped with the Tecan Fluent Robotic Liquid Handler and prepared for sequencing.

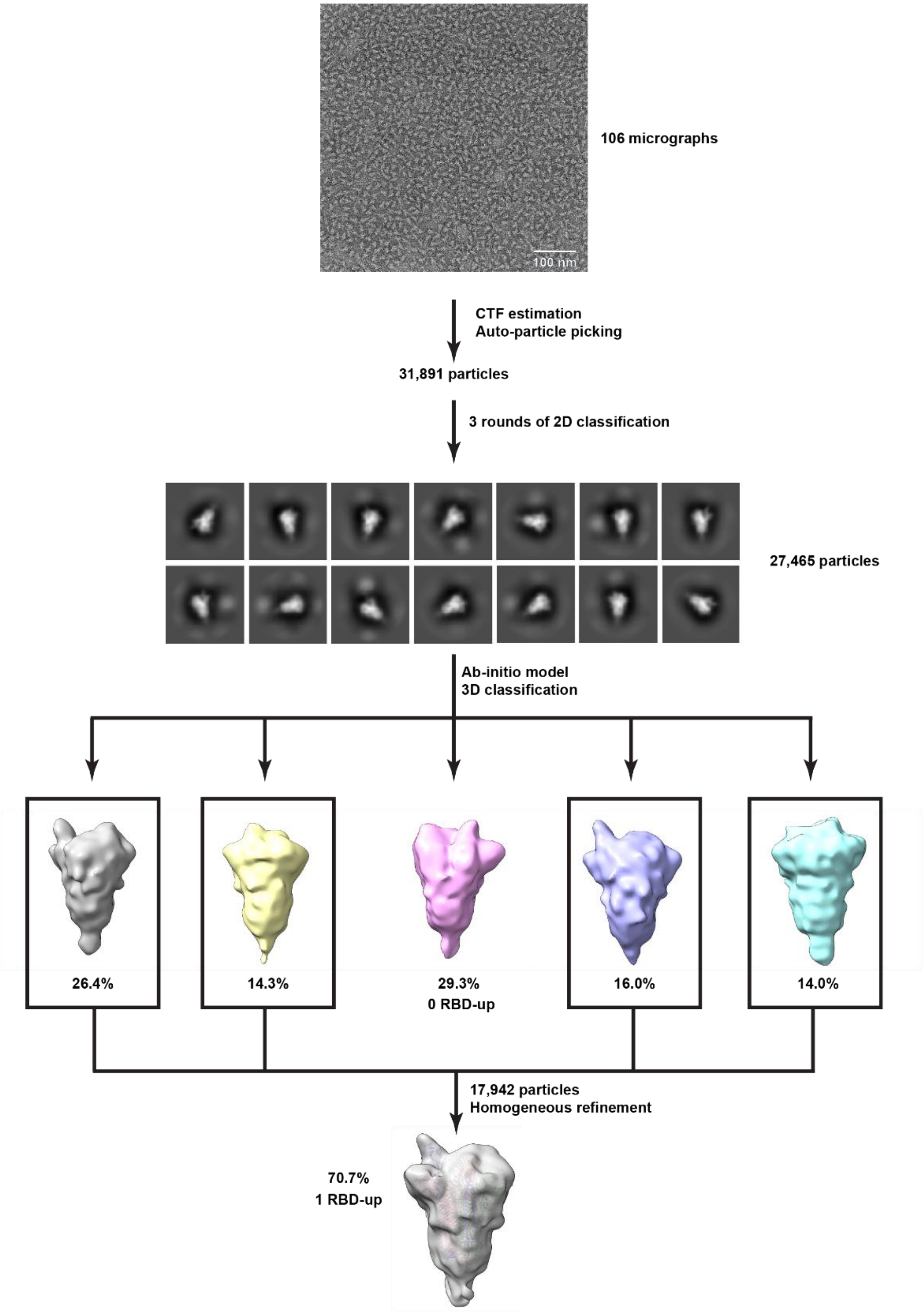


**Supplementary Fig. 3. Negative stain electron microscopy (nsEM) data processing workflow.** Schematic of pre-processing, classification, and refinement procedure to generate the map from 106 micrographs (see Methods). Boxed maps indicate the maps showing 1-RBD up EM density in alignment with HexaPro (S-6P) 1-RBD up structure (6XKL)[2].

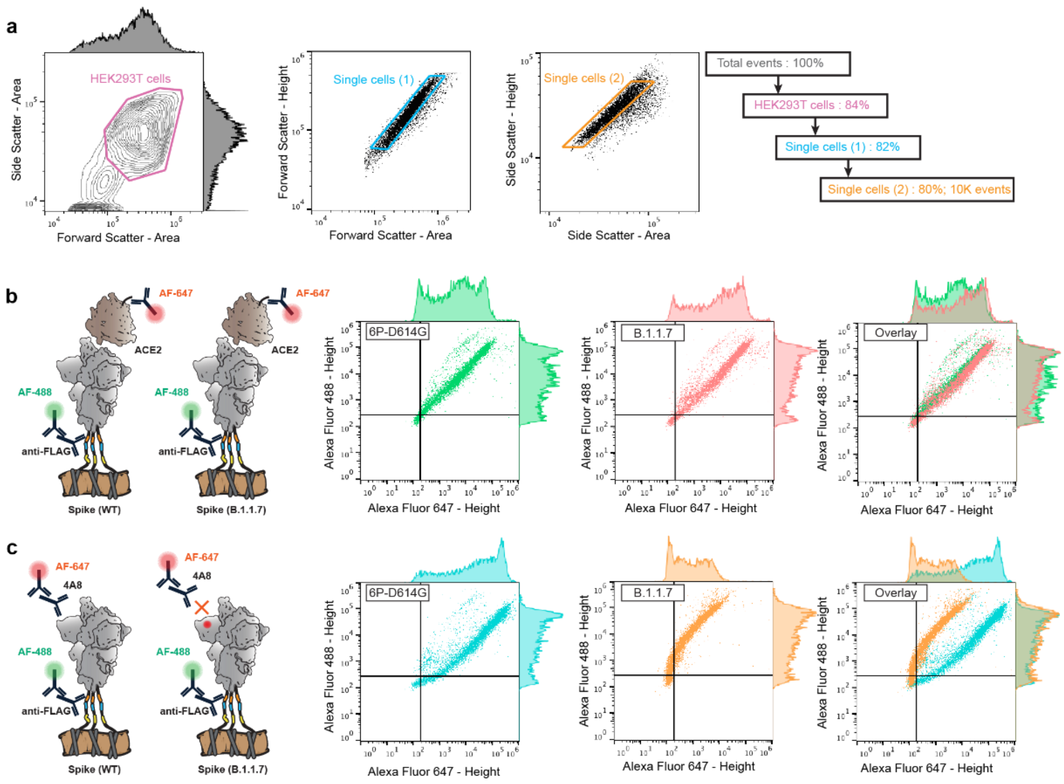


**Supplementary Fig. 4. Flow cytometry data processing.**
**(a)** Representative gating used to select for single HEK293T cells expressing surface displayed spikes. HEK293T cells identified with side scatter-area (SSA) vs forward scatter-area (FSA) gating followed by doublet discrimination using forward scatter-height (FSH) vs forward scatter-area (FSA) and side scatter-height (SSH) vs side scatter-area (SSA) gates. Representative cell% retained from each gating step shown on the right. **(b)** Graphical representation of the antibodies used (left) and the resulting flow cytometry data measuring ACE2 binding of 6P-D614G and B.1.1.7 spike variants (right). Single cells were analyzed using two fluorescent channels, Alexa Fluor-488 and Alexa Fluor-647 for measuring spike expression and antigenicity, respectively. **(c)** Graphical representation (left) and flow cytometry data measuring 4A8 antibody binding of 6P-D614G and B.1.1.7 spike variants (right). Histograms representing signal distributions for Alexa Fluor-488 and Alexa Fluor-647 are included for (b) and (c).

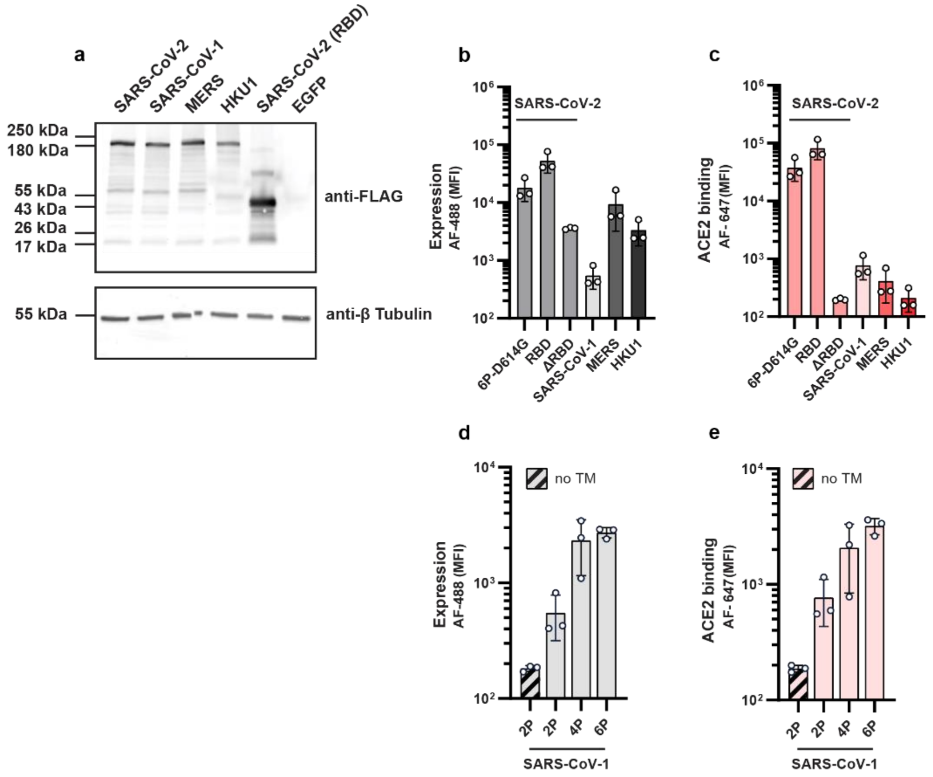


**Supplementary Fig. 5. Expression and ACE2 binding of diverse spike homologs. (a)** Western blot of spike displayed proteins and RBD from HEK293T cells. Western blots were stained with anti-FLAG antibodies. EGFP is included as a negative control for anti-FLAG, and anti-β tubulin used as a loading control. Raw expression **(b)** and ACE2 binding **(c)** signals measured via spike display and flow cytometry. The ΔRBD construct is a negative control for background ACE2 binding. Raw expression **(d)** and ACE2 binding **(e)** signals for pre-fusion stabilized SARS-CoV-1 using prolines. 2P = F799P + A874P. 4P = 2P + A881P + S924P. 6P = 4P + K968P + V969P. no TM = SARS-CoV-1 (2P) with no transmembrane domain, used as a negative control for background expression and ACE2 binding.
Circles represent biological replicates. Error bars = S.D.

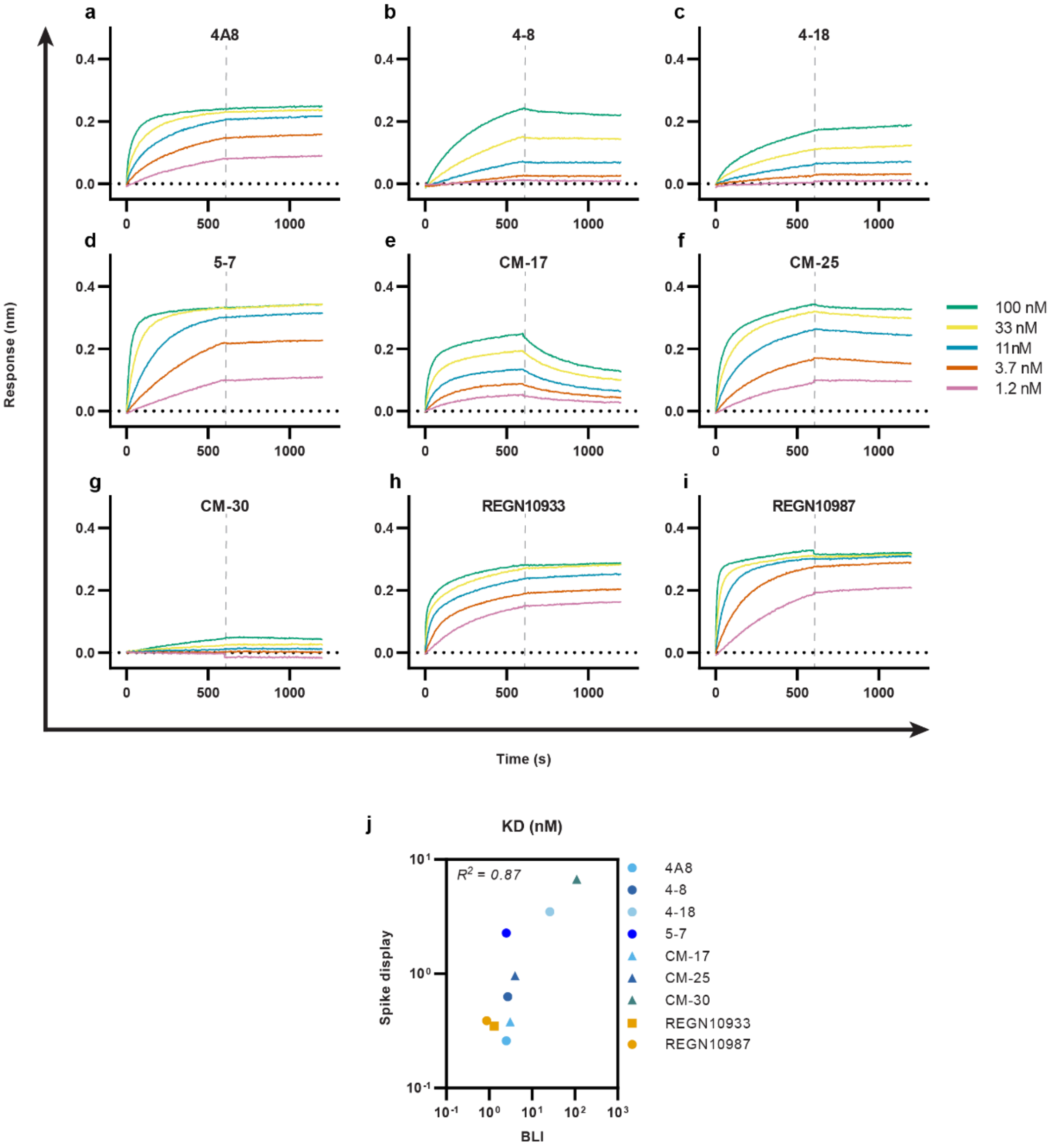


**Supplementary Fig. 6. Biolayer interferometry (BLI) of nAb binding to spike(6P-D614G).**

On- and off-rate curves for antibodies **(a)** 4A8, **(b)** 4-8, **(c)** 4-18, **(d)** 5-7, **(e)** CM-17, **(f)** CM-25, **(g)** CM-30, **(h)** REGN10933, and **(i)** REGN10987. **(j)** Correlation of the binding affinities measured by spike display and BLI.

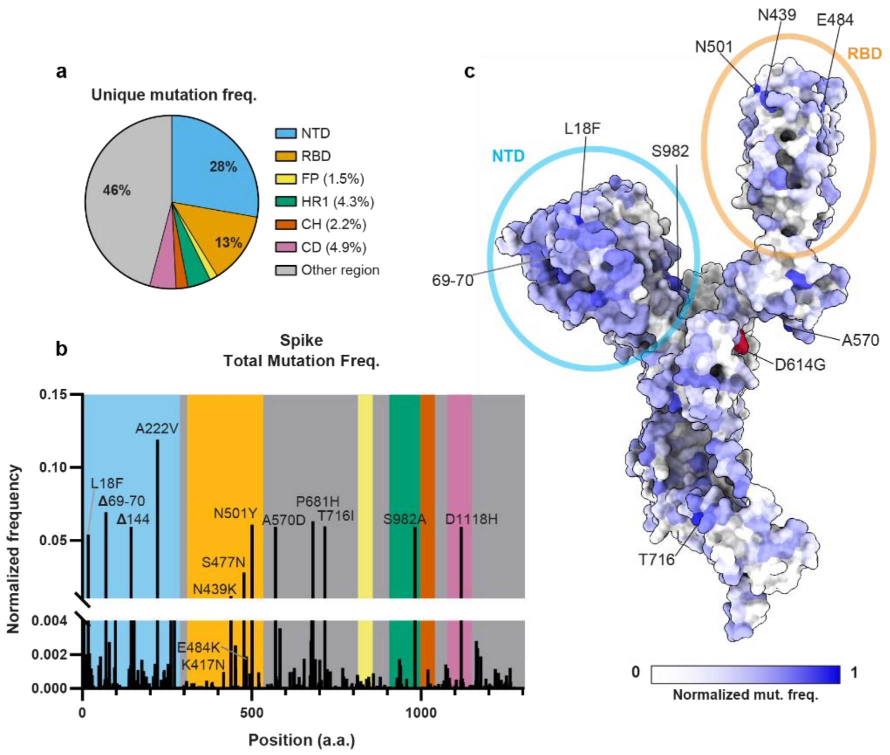


**Supplementary Fig. 7. Summary of SARS-CoV-2 spike mutation frequencies.**
**(a)** The distribution of spike mutations across key protein domains, as reported in the GISAID (accessed on 24/Feb/2021)[3,4]. **(b)** Normalized mutation frequencies, excluding the globally dominant D614G substitution. Spike domains are colored on the graph as in (a) and Fig. 2A. **(c)** A spike ectodomain monomer colored by mutation frequency (PDB: 7DDN[5]). White: invariant position (no mutations). A few key residues are labeled. Position 614, labeled in red, is excluded from the frequency calculations because it is now globally dominant[6].

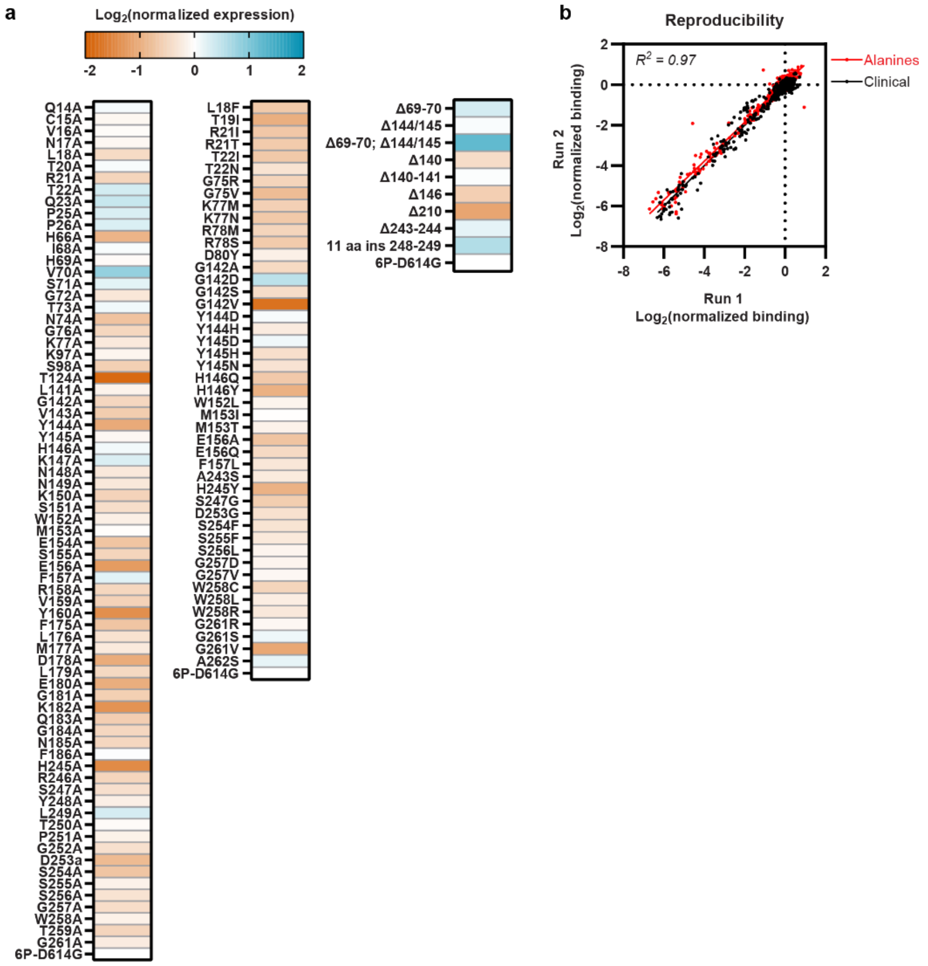


**Supplementary Fig. 8. Spike expression measurements and assay reproducibility.**
**(a)** Normalized spike expression for alanine and clinical variants in Figs. 2C, 3A, and 3E. **(b)** Comparison of the nAb binding data across two biological replicates and all spike variants indicates excellent reproducibility. Red: alanine scan; Black: all clinical variants. Pearson correlation was identical for both datasets.

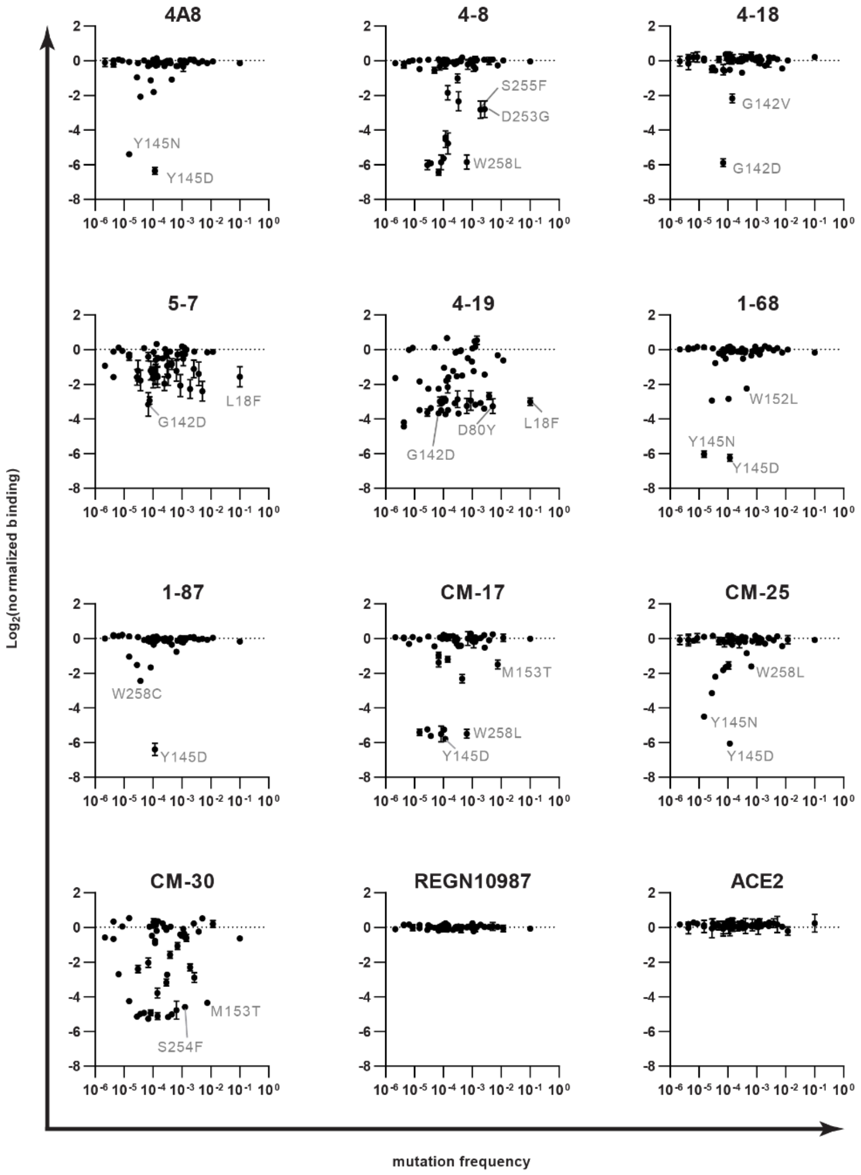


**Supplementary Fig. 9. Relationships between antibody binding and mutation frequency**
Comparison of mutation frequency as a function of NTD nAb escape outlined in Figure 3b. Mutations that decreased binding >16-fold typically appeared at very low frequencies (<0.001) in the GISAID. A few notable mutations are labeled. RGN10987 and ACE2 serve as RBD-binding negative controls; mutations in the NTD did not significantly affect their binding. All data points are a mean of at least two biological replicates. Error bars = S.D.

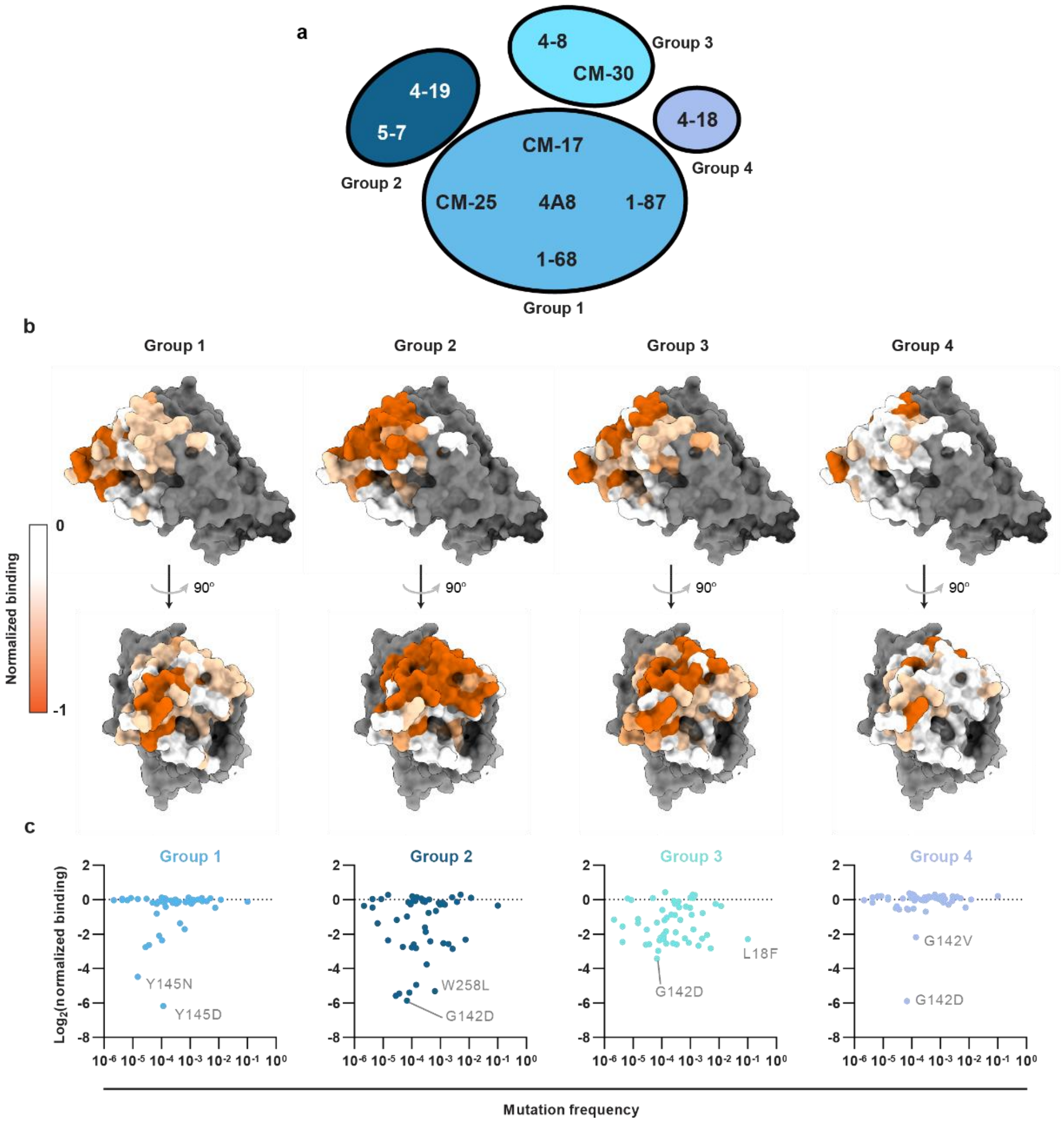


**Supplementary Fig. 10. Antibody classification and epitope mapping**
**(a)** NTD binding nAbs grouped according to Pearson correlation matrix (r > 0.5) from Fig. 3C. **(b)** Normalized binding data from figs. 2C and 3A were averaged for all nAbs in each group and normalized to a -1(loss of binding) to 0 (no change in binding) scale and superimposed on the NTD (PDB: 7DDN[5]). Gray: unmutated positions. Epitope maps highlight distinct binding patterns at the NTD supersite. **(c)** Comparison of mutation frequency as a function of NTD nAb escape for each group.

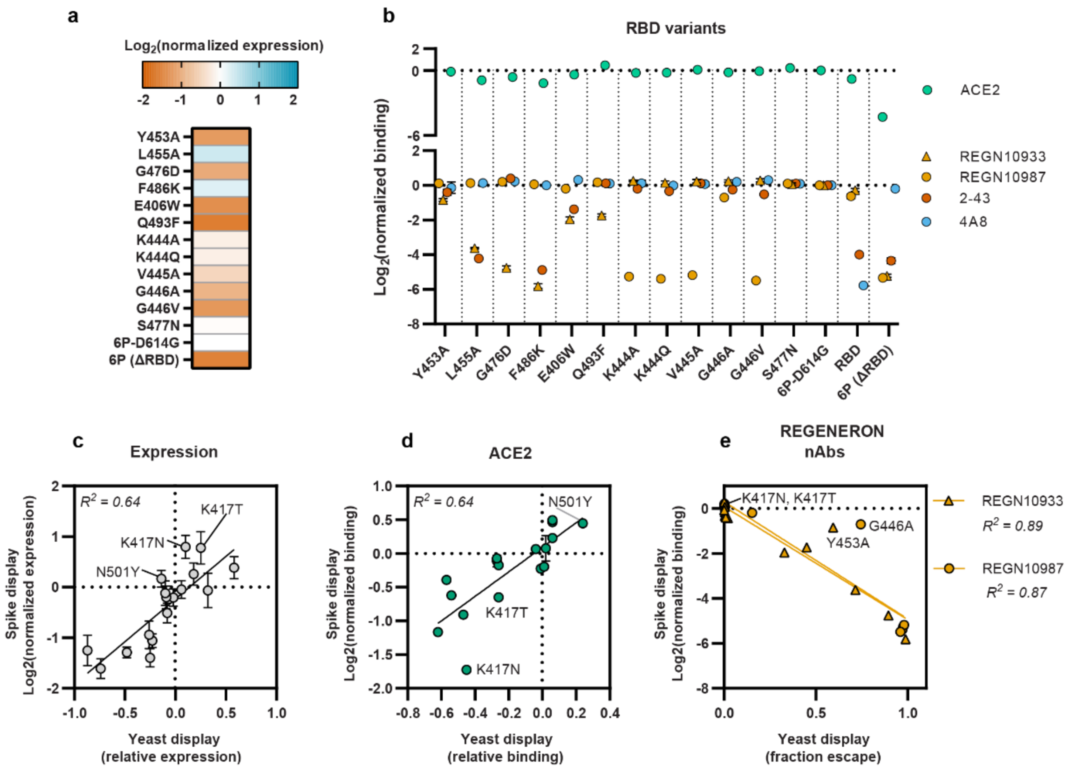


**Supplementary Fig. 11. Analysis of RBD mutations and comparison to yeast RBD display.** **(a)** Expression and **(b)** binding data for RBD variants that escape REGENERON nAbs REGN10933 and REGN10987. We also included the high prevalence S477N clinical mutation. Data in (a) and (b) were measured using spike display. (n = 3 biological replicates; error bars = S.D. Spike display correlates with yeast display of the isolated RBD[7,8] for **(c)** expression, **(d)** ACE2 binding, and **(e)** REGNERON nAb binding. Pearson correlation are indicated in each panel.

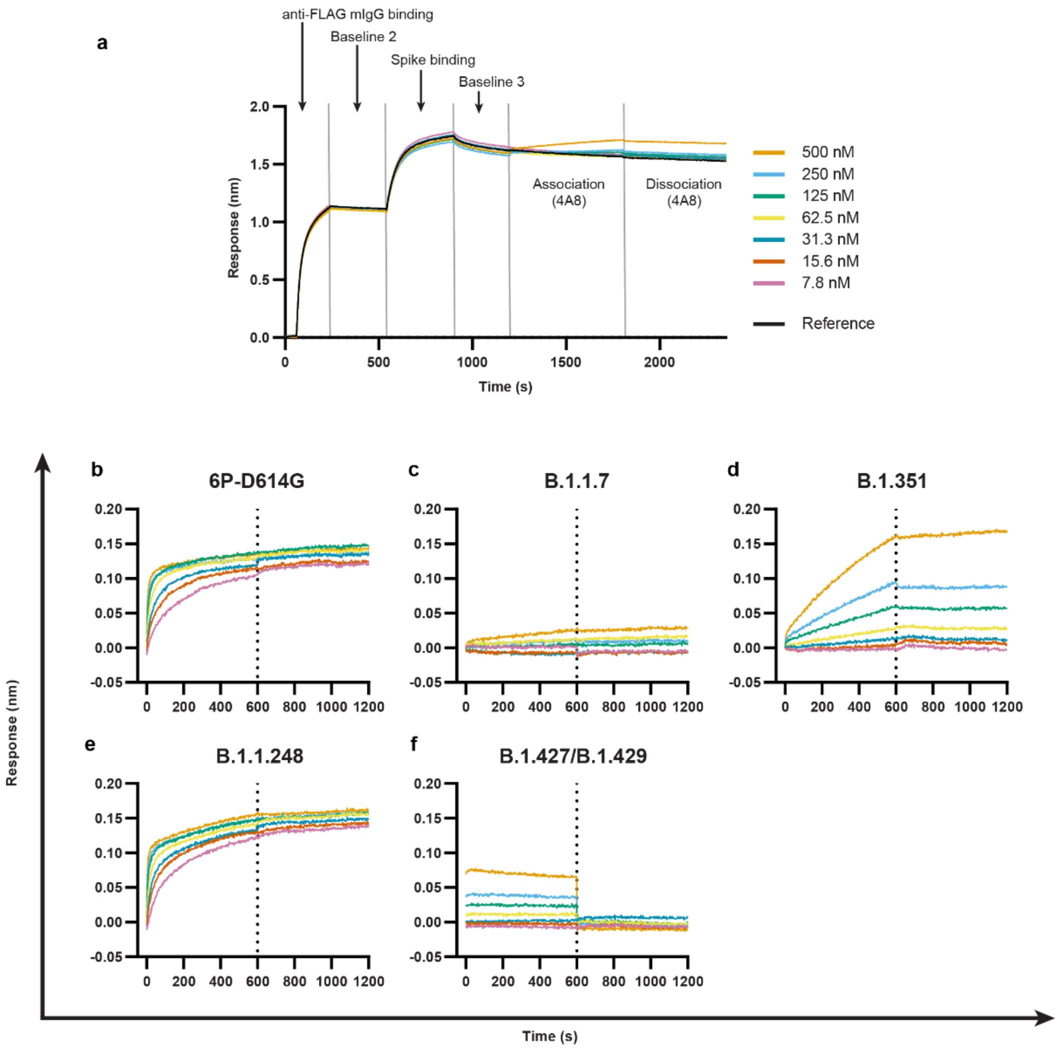


**Supplementary Fig. 12. Biolayer interferometry of VOC spikes cleaved from cell surfaces**

**(a)** BLI spectrogram showing the spike capture strategy. Anti-mouse Fc (AMC) tips are loaded with mouse anti-FLAG antibodies. The tips then capture SARS-CoV-2 spikes cleaved from cell surfaces with 3C protease (See Methods). BLI tips are further incubated in antibody (4A8) solutions of varying concentrations to observe "on" and "off" rates. (b-f) BLI on and off rate curves for 4A8 against 6P-D614G **(b),** B.1.1.7 **(c)**, B.1.351 **(d),** B.1.1.248 **(e)**, and B.1.427/B.1.429 **(f)** spikes. BLI experimental design matches that shown in (a). On/off rate transition for (b-f) indicated by vertical dotted lines.

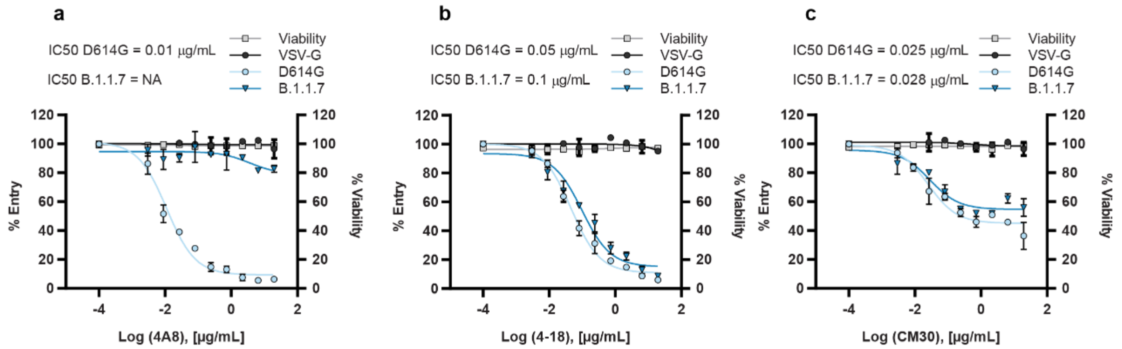


**Supplementary Fig. 13. Pseudovirus neutralization assays confirm B.1.1.7 partially escapes NTD antibodies.**

SARS-CoV-2 pseudovirus neutralization curves using 4A8 **(a)**, 4-18 **(b)**, and CM30 **(c)** nAbs. Results compare D614G (light blue) and B.1.1.7 (dark blue) pseudovirus variants (see Methods). VSV-G (black) included as a negative control for neutralization. Cell viability data included in gray. Dots and bars indicate mean and S.D of three replicates.

**Table S1. Comparison of antibody binding affinities using different platforms**

| Antibody | Specificity | Spike Display $K_D$ (nM) | BLI $K_D$ (nM)° | Literature $K_D$ (nM) |
|---|---|---|---|---|
| 4A8[9] | NTD | 0.26 ± 0.04 | 2.5 ± 0.06 | 0.996 ± 0.045[*] |
| 4-8[10] | NTD | 0.63 ± 0.04 | 2.7 ± 0.7 | 0.150 ± 0.037[#,§] |
| 4-18[10] | NTD | 3.5 ± 0.2 | 26 ± 4 | 0.216 ± 0.040[#,§] |
| 5-7[10] | NTD | 2.3 ± 0.1 | 2.5 ± 0.3 | 0.0950 ± 0.011[#,§] |
| CM-17[11] | NTD | 0.38 ± 0.04 | 3.1 ± 1 | 9.34 ± 0.16[#] |
| CM-25[11] | NTD | 0.96 ± 0.03 | 4.0 ± 0.2 | 9.44 ± 0.055[#] |
| CM-30[11] | NTD | 6.7 ± 0.6 | 110 ± 30 | 0.841 ± 0.84[#] |
| 2-43[10] | S1 | 0.22 ± 0.2 | N/A | 1.66 ± 1.5[&] |
| REGN10933[12] | RBD | 0.35 ± 0.05 | 1.3 ± 0.2 | 0.0417[@] |
| REGN10987[12]s | RBD | 0.39 ± 0.02 | 0.87 ± 0.07 | 0.0428[@] |

° Measured in this study using the full IgG.

[*] Measured via bio-layer interferometry (BLI).

[#] Measured via enzyme-linked immunosorbent assays (ELISAs)

[@] Measured via surface plasmon resonance (SPR).

[§] Digitized with WebPlotDigitizer and fit with a 4PL curve in GraphPad Prism 9. Error bar: 95% CI of the fit.

[&] Digitized with WebPlotDigitizer and fit with a 4PL curve in GraphPad Prism 9. Original data were measured via cell-surface competition binding assay. Error bar: 95% CI of the fit.

**Table S2. Biolayer interferometry of 4A8 to spike variants of concern.**

| Lineage | 4A8 mAb<br>$K_D$ (nM) |
|---|---|
| WT | 2.3 ± 0.4 |
| B.1.1.7 | N/D |
| B.1.351 | 34 ± 3 |
| B.1.1.248 | 2.0 ± 0.4 |
| B.1.427/B.1.429 | N/D |

N/D: Not determined because 4A8 binding was below the detection limit.
